# Local Delivery of Soluble Fractalkine (CX3CL1) Peptide Restore Ribbon Synapses After Noise-Induced Cochlear Synaptopathy

**DOI:** 10.1101/2024.02.20.581182

**Authors:** Vijayprakash Manickam, Sibaprasad Maity, Sree Varshini Murali, Dinesh Y. Gawande, Andrew R. Stothert, Lyudamila Batalkina, Astrid Cardona, Tejbeer Kaur

## Abstract

Efficacy of chemokine fractalkine isoforms was evaluated for restoration of loss of inner hair cell ribbon synapses and hearing after noise-induced cochlear synaptopathy (NICS). Previously, we have demonstrated a critical role for fractalkine signaling axis (CX_3_CL1-CX_3_CR1) in synaptic repair where in the presence of fractalkine receptor (CX_3_CR1) expressed by cochlear macrophages, the damaged synapses are spontaneously repaired. Here, we examined whether overexpression of fractalkine ligand (CX_3_CL1 or FKN) in the form of a peptide is effective in restoring the lost synapses and hearing after NICS. Remarkably, single transtympanic (TT) injection of soluble isoform of FKN (sFKN) peptide at 1 day after synaptopathic noise trauma showed significant recovery of ABR thresholds, ABR peak I amplitudes and ribbon synapses in both FKN-wildtype and knockout mice when compared to mice injected with full length membrane-bound FKN peptide (mFKN). Mechanistically, sFKN peptide treatment increased macrophage numbers in the cochlea and in the absence of those macrophages, sFKN failed to restore loss of synapses and hearing after NICS. Furthermore, sFKN treatment attenuated cochlear inflammation after noise overexposure without altering the expression of CX_3_CR1. Finally, sFKN peptide was detectable inside the cochlea localized to the sensory epithelium for 24 hours after TT injection. These data provide a robust proof-of-principle that local delivery of an immune factor, sFKN is effective in restoring lost ribbon synapses and hearing after NICS in a macrophage-dependent manner and highlights the potential of sFKN as an immunotherapy for cochlear synaptopathy due to noise or aging.

**Summary:** Transtympanic delivery of soluble FKN peptide is effective in restoring lost inner hair cell ribbon synapses and hearing after noise-induced cochlear synaptopathy in a macrophage-dependent manner.

## INTRODUCTION

Nearly 30 million Americans (18%) of ages 20-69 have hearing loss in both ears and 48 million have hearing loss in at least one ear from exposure to loud noise (National Institute on Deafness and Other Communication Disorders, NIDCD). The prevalence of noise-induced hearing loss in the military population is more significant than in the general public. As per the Department of Veterans Affairs, there are an estimated 1.3 million Veterans with a Service-related disability due to hearing loss (*1*). Noise exposure causes hearing loss due to damage to the sensory hair cells of the inner ear (*2*) or due to loss of synaptic contacts (a.k.a. ribbon synapse) between inner hair cells (IHCs) and peripheral dendrites of the spiral ganglion neurons (SGNs) (*3, 4*). Synaptic loss can trigger gradual degeneration of peripheral axons and ultimately death of SGNs (*4*). The consequences of synaptic and neuronal loss are deficits in hearing acuity, leading to difficulty in speech recognition and listening in noisy environments (*5*). This type of auditory dysfunction is referred to as noise-induced cochlear synaptopathy (NICS) or hidden hearing loss (HHL) because it can precede hair cell dysfunction or loss and is not readily diagnosed by standard clinical hearing tests such as auditory brainstem responses (ABRs) and distortion product otoacoustic emissions (DPOAEs). Notably, Veterans reporting high levels of military noise exposure display reduced suprathreshold ABR peak I amplitudes, which is a functional proxy read out for cochlear synaptopathy (*6*). Currently, there are no FDA-approved drugs that elicit regeneration of lost SGNs and restore their synaptic connections with the surviving hair cells. Moreover, loss of SGNs can limit the effectiveness of primary therapies for hearing loss such as hearing aids and cochlear implants or any future hair cell regeneration therapies. Neurotrophins such as NT-3, BDNF or agonists for Trk receptor (Neurotrophin receptor) can partially regenerate synapses in pre-clinical animal models of NICS (*7–11*). However, such partial effectiveness clearly expresses an urgent need to further delineate the cellular and molecular mechanisms of SGN survival and synapse repair or regeneration in order to identify newer targets and develop putative therapies to fully restore loss of synapses and hearing in NICS.

We have established a novel and critical role for cochlear macrophages (innate-immune cells) and fractalkine signaling in synaptic repair and SGN survival after NICS (*12, 13*). Fractalkine signaling axis (CX_3_CL1-CX_3_CR1) is a unique neuron-immune ligand-receptor pair in which fractalkine ligand (CX_3_CL1 or FKN), a chemokine, is constitutively expressed on neurons in the central nervous system (CNS) (*14, 15*) and on mouse SGNs and IHCs (*16*) and human SGNs (*17*). FKN is the exclusive ligand of a G-protein coupled receptor, CX_3_CR1, which is expressed by both human and mouse microglia (brain resident macrophages) (*18*) and cochlear resident macrophages (*19*). FKN is the only member of the CX3C, or delta, chemokine subfamily (*20, 21*). Unlike most other chemokine, FKN is a 395 amino acid (aa) type I transmembrane protein (*22*), including a signal sequence (aa 1-24), a chemokine domain (aa 25-100), a mucin stalk region (aa 101-336), a transmembrane segment (aa 337-357), and a cytoplasmic tail (aa 358-395). FKN exists in two different forms: a membrane-bound protein (mFKN) tethered to neuronal membranes by a mucin-like stalk and a soluble factor (sFKN) released upon cleavage of its N-terminal chemokine domain by metalloproteinases (*23, 24*). The soluble chemokine domain of fractalkine acts as a chemoattractant promoting migration of immune cells expressing CX_3_CR1, while the membrane-tethered mucin-stalk of fractalkine acts as an adhesion molecule for leukocytes to endothelium during tissue injury (*25, 26*). Pre-clinical animal studies in the CNS have shown that neurotransmitter glutamate-induced excitotoxic injury activate and recruit microglia to the site of injury and that those microglia protect neurons and improve synaptic recovery following excitotoxic damage (*27–31*). Such microglia-mediated protection against CNS excitotoxicity have been partly attributed to fractalkine signaling (*32–42*). Moreover, overexpression of the soluble isoform of FKN (sFKN) confers neuroprotection, synaptic recovery, and improved function in animal models of several neurodegenerative and neuroinflammatory disorders (*43–56*).

We have demonstrated that noise trauma that induces a temporary shift in hearing thresholds (TTS) without any significant hair cell damage and/or loss causes rapid degeneration of IHC ribbon synapses and migration of resident cochlear macrophages into the damaged IHC-synaptic region (*12*). We and others have reported that noise-damaged IHC synapses can undergo spontaneous repair/regeneration in zebrafish, mice, guineapigs and gerbil (*12, 13, 57–66*). We have also shown that genetic disruption of fractalkine signalling due to lack of CX_3_CR1 receptor on macrophages or absence of cochlear resident macrophages hampers spontaneous synaptic repair and augments SGN loss and inflammation after noise trauma (*12, 13, 67*). These results suggest that endogenous intact fractalkine signaling axis plays a key role in synaptic repair and SGN survival through suppression of inflammation in noise-damaged cochlea. So, we probed, does boosting the cochlear FKN levels adequate to restore loss of synapses and hearing after NICS? Our data serves as a robust proof-of-principle that local delivery of an immune factor, sFKN peptide is effective in restoring the loss of IHC ribbon synapses and hearing after NICS and such sFKN-mediated synaptic repair is dependent on cochlear macrophages. Furthermore, our data demonstrate that transtympanically delivered sFKN peptide localizes to the sensory epithelium and resolves inflammation due to NICS. These findings highlight the potential of sFKN as an immunotherapy for cochlear synaptopathy due to either noise or biological aging.

## RESULTS

### sFKN is more effective than mFKN in recovering ABR threshold shifts after NICS in FKN KO mice

Our overarching goal in this study was to evaluate the efficacy of FKN isoforms (membrane-bound and soluble) in restoration of loss of IHC ribbon synapses and hearing after NICS. We first assessed the efficacy of FKN isoforms in the recovery of ABR threshold shifts and peak I amplitudes after NICS. Gross hearing function and IHC ribbon synapse density in unexposed FKN KO mice were comparable to the age-matched unexposed WT mice (Fig. S1). At 1 DPNE, both FKN WT and KO mice showed similar degree of elevation in ABR thresholds with FKN KO mice displaying ∼10 dB higher threshold shifts than FKN WT mice when compared to the thresholds before noise exposure (Fig. 1B and C). There was an average 40 dB and 50 dB shift in ABR thresholds at stimulus frequencies of 16 kHz and above at 1 DPNE in the FKN WT and KO mice, respectively. By 15 DPNE, vehicle-treated WT mice showed a robust recovery in ABR threshold shift whereas, similar level of recovery was not observed in the vehicle-treated FKN KO mice (Fig. 1B and C). Remarkably, this loss of function in the noise-exposed FKN KO mice was significantly rescued by the single transtympanic administration of sFKN peptide, and the degree of recovery was comparable to vehicle-treated FKN WT mice (Fig. 1F). Whereas there was little to no recovery in ABR threshold shifts in the noise-exposed FKN KO mice injected with either mFKN peptide (Fig. 1E) or control peptide (Fig. 1D). There was no significant difference in the DPOAE levels at 15 DPNE when compared to pre-noise exposure among the experimental groups (Fig. S2). Furthermore, lack of FKN did not influence the survival of inner and outer hair cells in both unexposed and noise-exposed mice (Fig. S3).

**Fig. 1.**
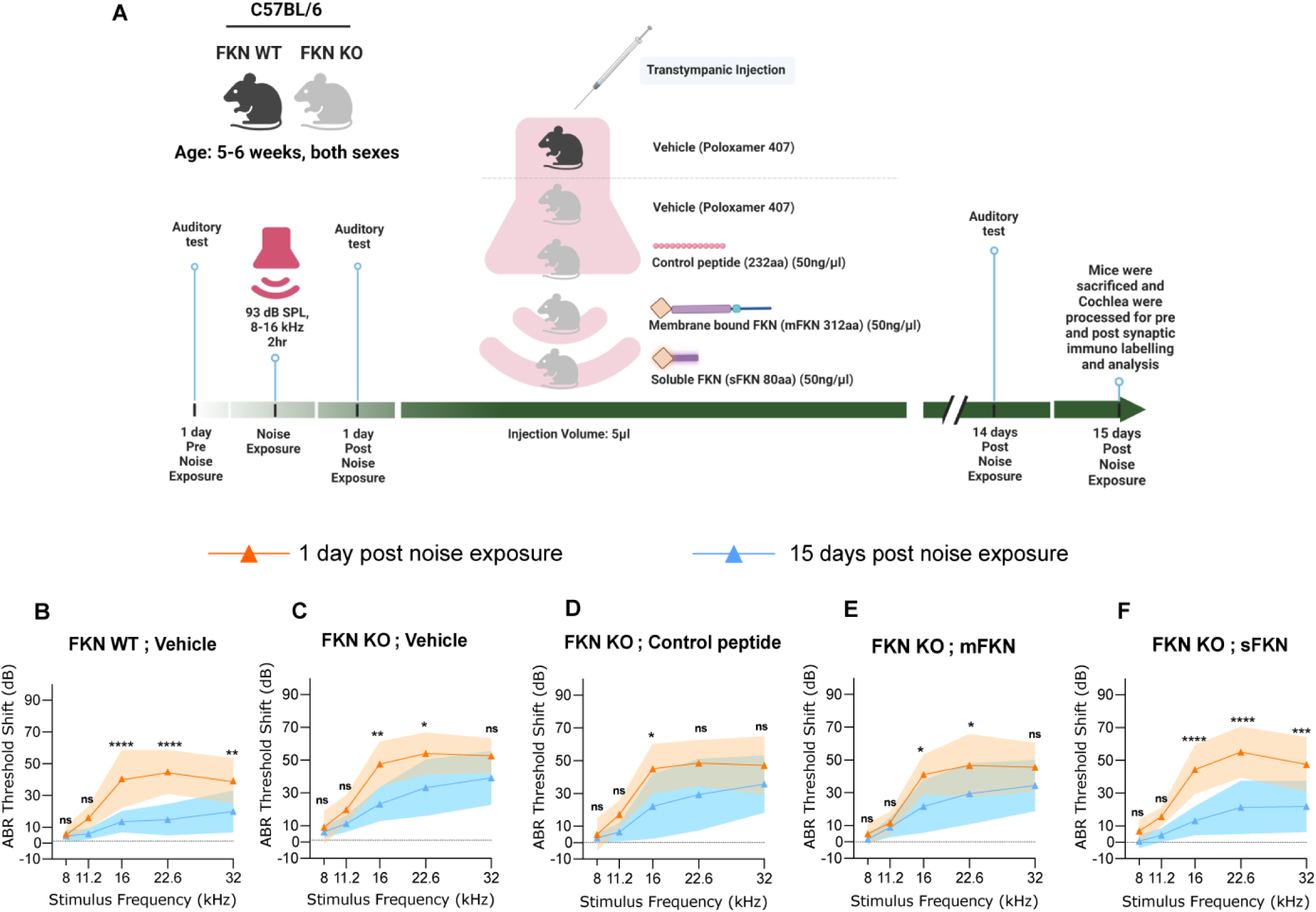
ABR threshold shifts at 1 DPNE and 15 DPNE. (A) Study design created with BioRender.com. (B-F) ABR threshold shifts at 1 DPNE and 15 DPNE for 2 hours at 93 dB SPL at 8-16 kHz octave band from (B) FKN WT mice treated with vehicle (N=8), and FKN KO mice treated with (C) vehicle (N=6) (D) control peptide (N=7) (E) membrane-bound FKN peptide (mFKN) (N=9) and (F) soluble FKN peptide (sFKN) (N=8). Values are means ± SD. *P < 0.05, **P < 0.01, ***P < 0.001, ****P < 0.0001 and ns, non-significant at respective stimulus frequency. *Represents comparison between 1 DPNE and 15 DPNE, 2-way ANOVA, Sidak’s multiple comparisons. Dashed line represent threshold shifts prior to noise exposure (baseline).

### sFKN is more effective than mFKN in restoring ABR peak I amplitudes after NICS in FKN KO mice

Despite of the spontaneous recovery of ABR threshold shifts by 15 DPNE, FKN WT mice have not recovered the reduced ABR-P-I amplitude at 32 kHz (basal cochlear region) (Fig. 2A), displaying the hallmark of noise-induced hidden hearing loss (*4*). The functional input/output series shows significant reduction in ABR P-I amplitudes at suprathreshold sound levels at 1 DPNE and 15 DPNE compared to amplitudes at pre-noise exposure (baseline). Similarly, none of the FKN KO mice injected with either vehicle, control peptide or mFKN peptide showed recovery in reduced ABR-P-I amplitude at 32 kHz (Fig. 2B, C and D). However, FKN KO mice TT injected with a single dose of sFKN peptide displayed robust recovery of reduced ABR-P-I amplitudes to baseline amplitudes at 15 DPNE (Fig. 2E). There was no significant change observed in ABR-P-I amplitudes at 8 kHz and 16 kHz in any of the experimental groups (Fig. S4). Similarly, ABR-P-I latencies at 8 kHz, 16 kHz and 32 kHz were largely unaffected among the experimental groups at all recovery time points after noise trauma (Fig. S5). Among the FKN isoforms, overexpression of sFKN peptide restored ABR thresholds and P-I amplitudes in both male and female FKN KO mice but the recovery was more robust in female FKN KO mice (Fig. S6). Collectively, this and above data implicate that cochlear endogenous FKN regulate the recovery of loss of hearing sensitivity and that overexpression of sFKN peptide in mice lacking endogenous FKN is more effective than mFKN peptide in restoring hearing loss after NICS.

**Fig. 2.**
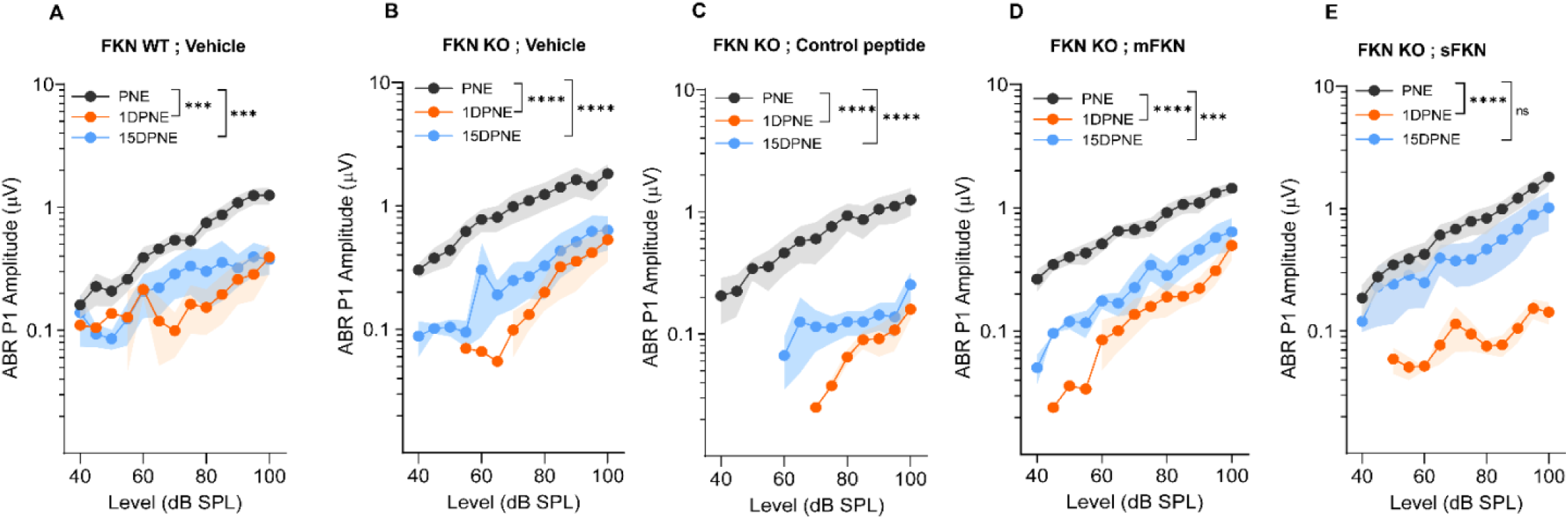
ABR peak I amplitude at 32 kHz at Pre-noise exposure (PNE), 1 DPNE and 15 DPNE. ABR peak I amplitudes at 32 kHz in (A) FKN WT mice treated with vehicle (N=8), and in FKN KO mice treated with (B) vehicle (N=6) (C) control peptide (N=7) (D) membrane-bound FKN peptide (mFKN) (N=10) and (E) soluble FKN peptide (sFKN) (N=8). Values are means ± SD. ***P < 0.001, ****P < 0.0001 and ns, non-significant. *Represents the comparison between the experimental time points as indicated with parenthesis. 1-way ANOVA, Tukey’s multiple comparisons test.

### Single TT injection of sFKN peptide restore noise-damaged ribbon synapses in FKN KO mice

In order to validate the functional recovery of the IHC ribbon synapses, microdissected cochleae were immunolabelled for presynaptic CtBP2 ribbons and postsynaptic GluA2 AMPA receptors (Fig. 3A). Compared to unexposed groups, there was a significant reduction in the absolute CtBP2 punctae (Fig. 3B), absolute GluA2 punctae (Fig. 3C) and paired synapses (Fig. 3D) per IHC (from ∼18 to ∼10 punctae or synapses per IHC) in the basal region of the cochlea of noise-exposed groups at 15 DPNE, irrespective of the genotype. Among the noise-exposed FKN KO groups, mice TT injected with sFKN peptide showed a significant restoration of GluA2 punctae and paired synapses per IHC (from ∼10 to ∼17 punctae or synapses per IHC) when compared to mice injected with mFKN peptide or with control peptide (Fig. 3C and D). Notably, the synaptic repair due to the sFKN peptide treatment is comparable to the IHC synaptic density in unexposed FKN KO group. Such structural recovery of synapses positively correlates with the functional recovery of ABR peak I amplitude (Fig. 2E) and suggest that between the two FKN isoforms, sFKN is more efficacious in restoring damaged synapses after NICS, both structurally and functionally. There was no significant loss in the absolute CtBP2 punctae, GluA2 punctae and paired ribbon synapses per IHC in apical and middle cochlear regions in all the noise-exposed experimental groups at 15 DPNE (Fig. S7).

**Fig. 3.**
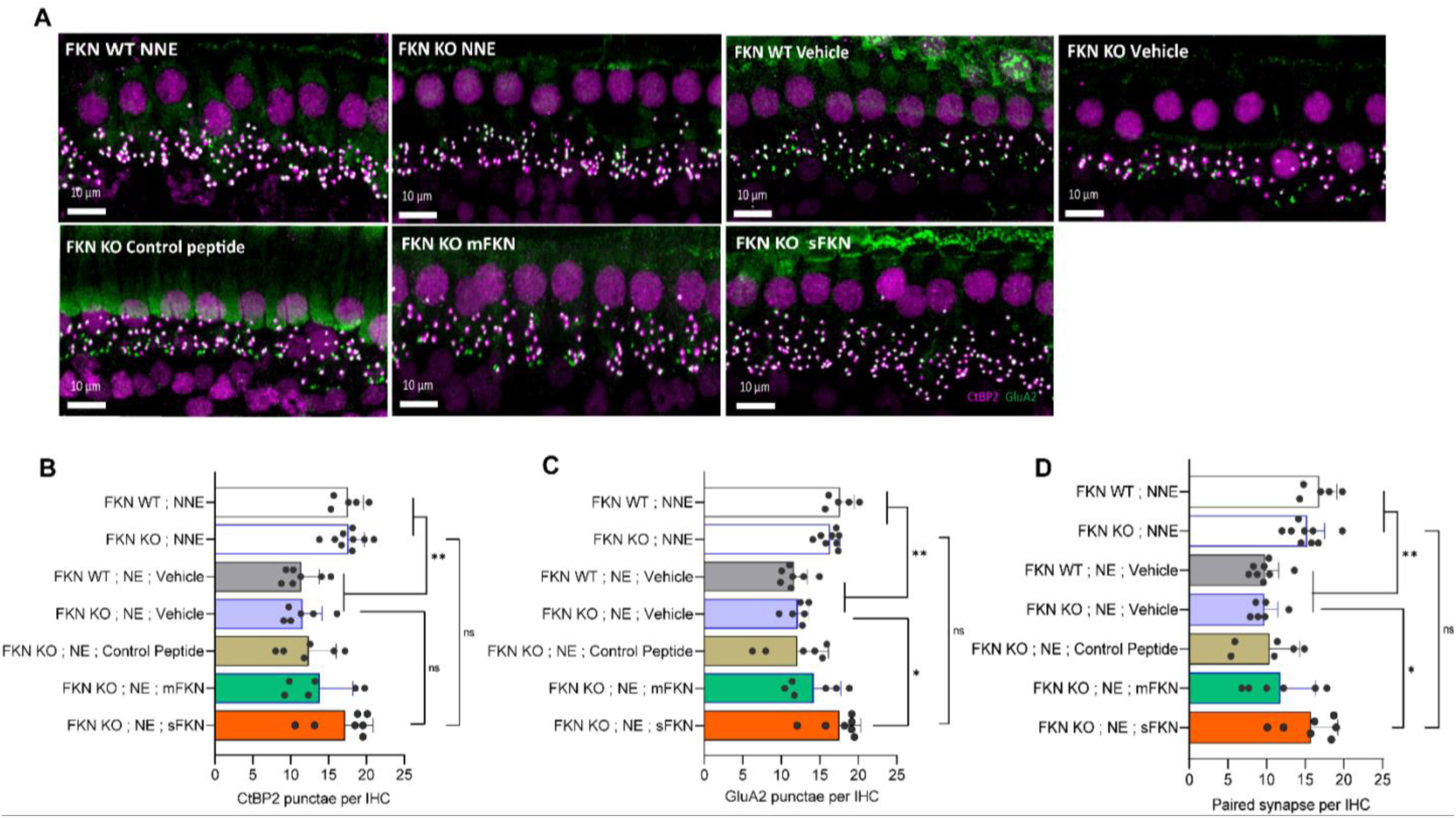
Inner hair cell ribbon synapses in basal cochlear region at 15 DPNE. (A) Representative micrographs showing IHC paired ribbon synapses in basal cochlear region after 15 days of synaptopathic noise exposure. (B) Absolute CtBP2 punctae per IHC. (C) Absolute GluA2 punctae per IHC. (D) Paired ribbon synapses per IHC. Values are mean ± SD. Each dot in the graphs represents a mouse. **P < 0.01 between no noise exposed (NNE) vs. noise exposed vehicle treated FKN WT and FKN KO mice; *P < 0.05 between noise exposed FKN KO mice treated with vehicle or sFKN peptide; ns: non-significant between NNE and NE sFKN treated FKN KO mice. 1-way ANOVA, Dunnett’s multiple comparisons test. N=5-9 mice per experimental group.

### Single TT injection of sFKN peptide restore noise-damaged ribbon synapses in FKN WT mice

As there was no spontaneous synaptic repair observed in the basal cochlear region of the noise-exposed FKN WT mice at 15 DPNE (Fig. 3) therefore, we hypothesized that this is likely due to the inadequate levels of cochlear endogenous FKN ligand promoting synaptic repair after NICS (Fig. S8). To test whether sFKN peptide also restores noise-damaged ribbon synapses in FKN WT mice, we overexpressed sFKN peptide by TT injection in FKN WT mice at 1 day post noise exposure and analyzed ABR P-I amplitudes at 32 kHz (Fig. 4A) and IHC ribbon synapse density in the basal cochlear region (Fig. 4B and C). There was a significant reduction in ABR P-I amplitude at 1 DPNE compared to the pre-noise exposure (baseline) that was partially but significantly recovered by 15 DPNE in FKN WT mice due to sFKN peptide treatment post noise exposure. Moreover, noise-damaged IHC ribbon synapses were robustly restored as a result of treatment with sFKN peptide (from ∼ 9 to ∼13 synapses per IHC). This data suggests that the overexpression of sFKN peptide by single TT injection can restore noise-damaged IHC ribbon synapses both functionally and structurally in the wild-type mice besides FKN KO mice.

**Fig. 4.**
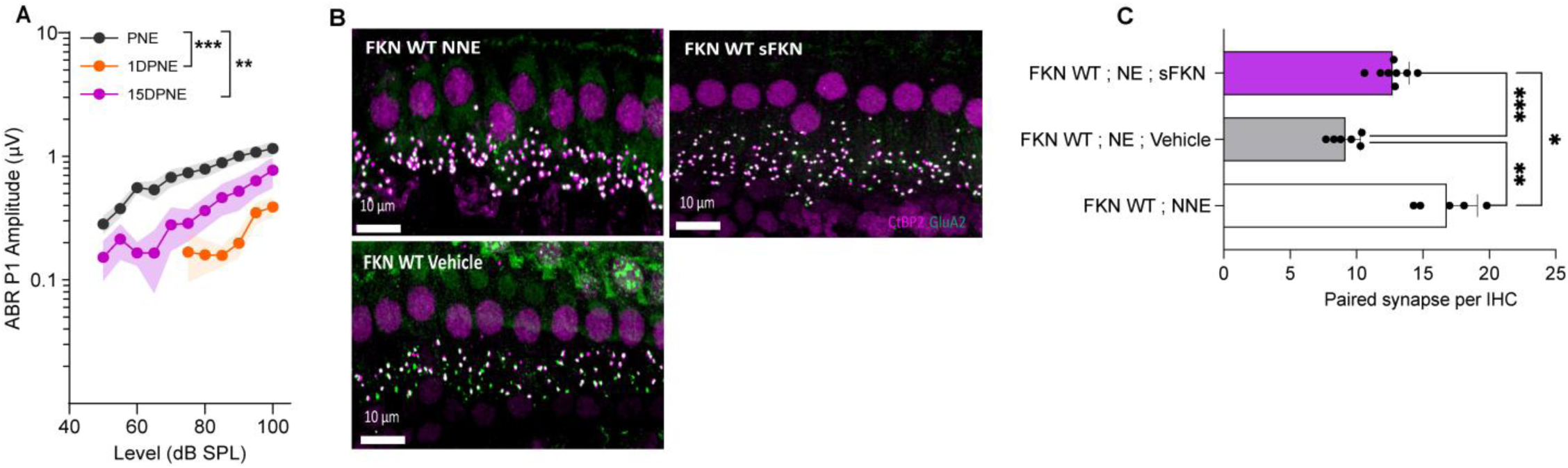
ABR peak I amplitude and IHC ribbon synapse density in FKN WT mice TT injected with sFKN peptide. (A) ABR peak I amplitude at 32 kHz at pre-noise exposure (PNE), 1 DPNE and 15 DPNE of FKN WT mice (N=8) injected TT with a single dose of sFKN peptide. (B) Representative micrographs showing IHC ribbon synapses from basal cochlear region at 15 DPNE. (C) Paired ribbon synapses per IHC. Values are means ± SD. Each dot in the graph represents a mouse. *P < 0.05, **P < 0.01, ***P < 0.001 and ns, non-significant. 1-way ANOVA, Tukey’s multiple comparisons test. FKN WT NNE and FKN WT vehicle data shown in B and C have been reused from Fig. 3 in order to reduce the use of numbers of mice as per IACUC policies.

### Cochlear macrophages are indispensable for sFKN-mediated IHC synaptic repair after NICS

We have previously reported that cochlear resident macrophages are activated and migrate immediately into the sensory epithelium in close proximity to the damaged IHC synaptic region following a synaptopathic noise trauma (*12*). In order to determine whether chemokine FKN influence macrophage localization and numbers in the damaged sensory epithelium after NICS, cochlear whole mounts were immunolabeled for CD45, a pan-leukocyte marker. The macrophage numbers significantly increased in the middle (∼2.5-fold) and basal (∼2-fold) cochlear region of the damaged sensory epithelium of FKN WT mice after noise trauma nevertheless, such an increase in macrophage numbers was not observed in the noise-damaged sensory epithelium of FKN KO mice (Fig. 5A and B). There was no significant change in macrophage numbers in the apical region of the cochlea in both noise-exposed FKN WT and KO groups when compared to unexposed mice. This data suggests that chemokine FKN which is constitutively expressed by IHCs and SGNs control macrophage numbers in the sensory epithelium in response to NICS.

**Fig. 5.**
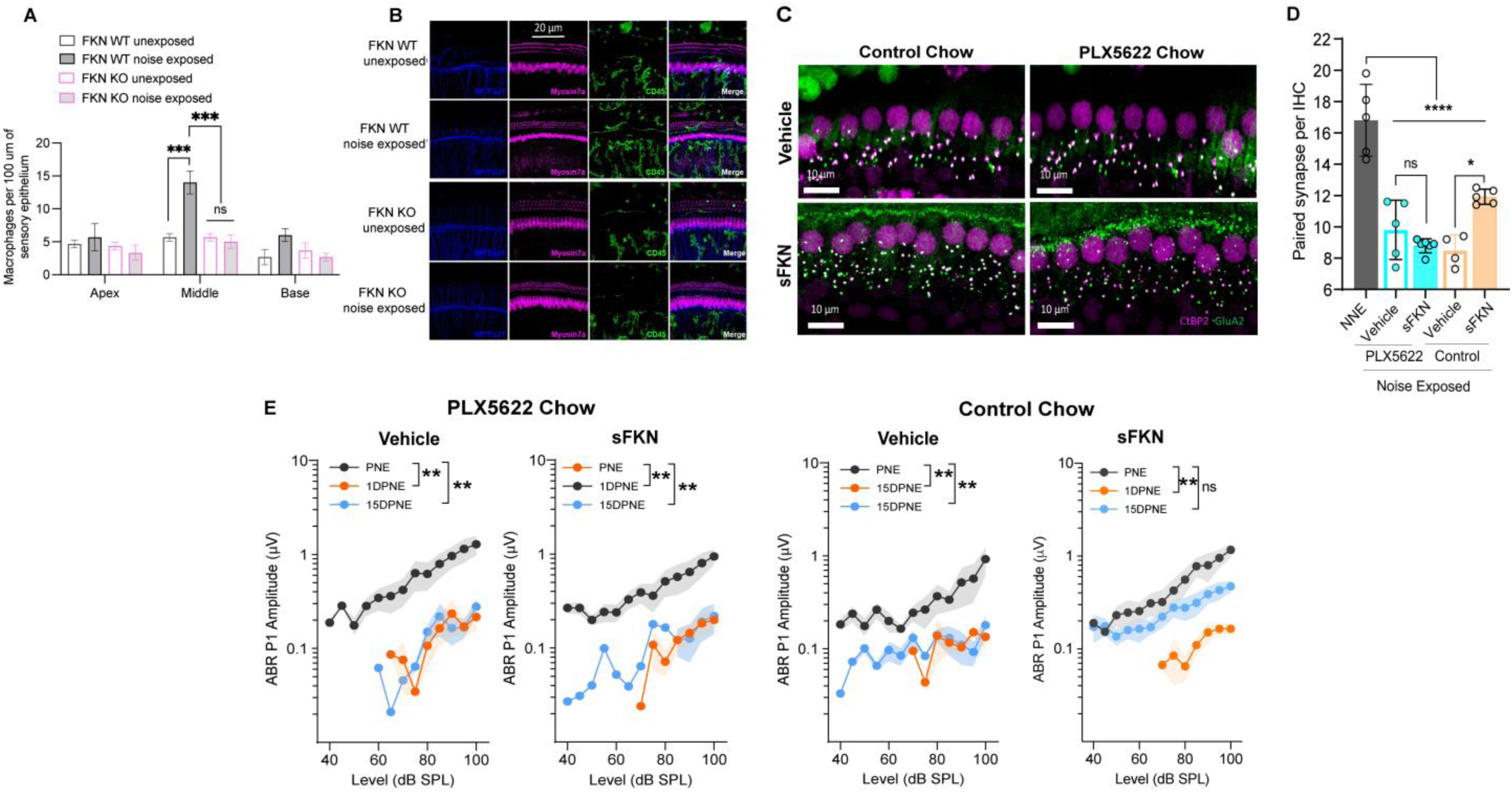
sFKN fails to restore noise-damaged IHC ribbon synapses and ABR Peak I amplitudes in FKN WT mice lacking cochlear resident macrophages. (A) Macrophage density in the sensory epithelium of apex, middle and basal cochlear regions of unexposed and noise-exposed FKN WT and FKN KO mice. 2-way ANOVA, Tukey’s comparison test. N=3 mice per genotype. (B) Representative micrographs showing CD45-immunolabeled macrophages in the sensory epithelium of the middle cochlear region of unexposed and noise-exposed FKN WT and FKN KO mice. (C) Representative micrographs showing IHC paired ribbon synapses from basal cochlear region of vehicle- or sFKN peptide-treated FKN WT mice in the presence (control chow) or absence (PLX5622 chow) of macrophages at 15 DPNE. (D) Quantification of paired ribbon synapses per IHC in vehicle- or sFKN peptide-treated FKN WT mice in the presence (control chow) or absence (PLX5622 chow) of macrophages at 15 DPNE. Gray bar represents unexposed (NNE) FKN WT mice data reused from Fig. 4C for comparison purposes and to reduce the use of numbers of mice as per IACUC policies. 2-way ANOVA, Tukey’s comparison test. (E) ABR Peak I amplitudes at 32 kHz in vehicle- or sFKN peptide-treated FKN WT mice in the presence (control chow) or absence (PLX5622 chow) of macrophages at 15 DPNE. 1-way ANOVA, Dunnett’s multiple comparisons test. N=5-7 mice per experimental group (D and E). Values are means ± SD. *P < 0.05, **P < 0.01, ***P < 0.001, ****P < 0.0001 and ns, non-significant.

FKN binds to it sole receptor, CX_3_CR1 which is thought to be specifically expressed by macrophages (*19*). This fact and the data that both FKN (Fig. 5A and B) and CX_3_CR1 (*16, 67*) regulates macrophage density in the damaged cochlea prompted us to ask whether the sFKN-mediated synaptic repair is a direct effect of sFKN on IHC ribbon synapses and SGNs or does it require cochlear macrophages expressing CX_3_CR1. To determine whether macrophages are necessary for sFKN-mediated synaptic repair after NICS, cochlear macrophages were depleted in FKN WT mice via treatment with CSF1-R inhibitor, PLX5622 chow for 2 weeks which effectively results in ∼94% ablation of cochlear resident macrophages (*13*) (also see Fig. S9). Following macrophage ablation, noise-exposed FKN WT mice were TT injected with either vehicle or sFKN peptide and synapses and function were analyzed at 15 DPNE. Depletion of cochlear macrophages with PLX5622 abrogated the ability of sFKN peptide to restore noise-damaged ribbon synapses (Fig. 5C and D) and ABR Peak I amplitudes (Fig. 5E) in FKN WT mice. Whereas sFKN peptide restored the loss of synapses and function in the presence of cochlear resident macrophages (control chow) in FKN WT mice (Fig. 5C-E, also see Fig. 4). The present data also corroborates our previous report by Manickam et al., 2023 (*13*) that the absence of cochlear macrophages impairs the recovery of ABR thresholds shifts which is not affected by sFKN peptide treatment, whereas elevated ABR thresholds recover in the presence of macrophages with or without sFKN treatment (Fig. S10A-D). DPOAE levels were comparable among the experimental groups (Fig. S10E-H). Mechanistically, these data implicate that sFKN peptide restore the noise-induced loss of IHC ribbon synapses and function in a macrophage-dependent manner.

### NICS-induced proinflammatory cytokines is diminished by sFKN

Noise-induced cochlear synaptopathy is associated with activation of macrophages (*12*) and upregulation of inflammation (*68*). Accordingly, as CX_3_CL1 signaling has been shown to attenuate the production of pro-inflammatory cytokines and decrease activation of macrophages *in vitro* and *in vivo* following various inflammatory stimuli (*14, 69–76*), we examined the pro-inflammatory reducing and anti-inflammatory enhancing properties of sFKN in response to NICS in the FKN WT mice. Cochlear protein lysates from no-noise exposed (NNE), noise-exposed plus TT injection of vehicle at 1 day post exposure (NE; vehicle) and noise exposed plus TT injection of sFKN peptide at 1 day post exposure (NE; sFKN) were collected at ∼48 hours post noise exposure and analyzed for cytokines levels using Luminex-based multiplex technique. Production of pro-inflammatory cytokines such as IFN ß, IL-2, IL-6, and IL-23 were significantly elevated in the cochleae injected with vehicle at 1 day after a synaptopathic noise trauma when compared to no-noise exposed FKN WT mice. However, transtympanic treatment with sFKN peptide attenuated or maintained the levels of these cytokines similar to the levels found in the no-noise exposed FKN WT mice (Fig. 6A-D). Similarly, levels of anti-inflammatory cytokines such as IL-10, IL-22 and IL-33 were significantly reduced in the cochleae injected with vehicle at 1 day after a synaptopathic noise trauma when compared to no-noise exposed FKN WT mice but transtympanic treatment with sFKN peptide after noise trauma elevated or maintained their levels close to baseline (Fig 6E-G). We also measured the messenger RNA (mRNA) levels of cochlear CX_3_CR1 receptor that is exclusively expressed by macrophages in FKN WT mice following TT injection of sFKN peptide by RT-qPCR. No substantial change was detected in the expression of CX_3_CR1 mRNA among the unexposed, noise-exposed vehicle injected and noise-exposed sFKN injected groups at 2 and 24 hours post sFKN peptide injection (Fig. S11). Collectively, these results suggest that sFKN-mediated IHC synaptic repair after NICS is likely attributed to modulation of cochlear inflammation in response to NICS and not due to changes in the expression of CX_3_CR1 receptor on cochlear macrophages.

**Fig. 6.**
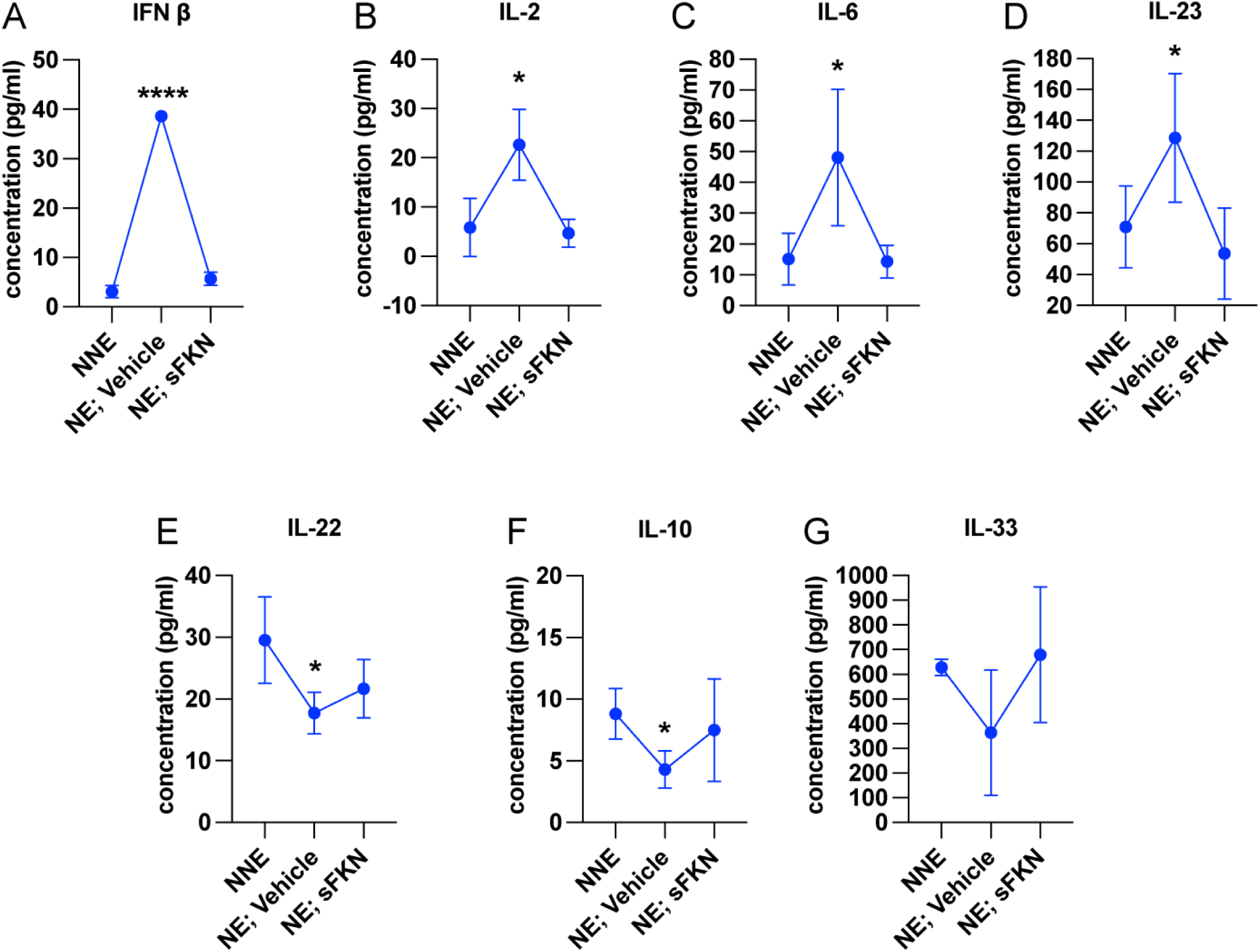
sFKN peptide regulates cochlear inflammation profile after NICS. Luminex assay-based quantification of the levels of cytokines (A) IFN ß, (B) IL-2, (C) IL-6, (D) IL-23, (E) IL-22, (F) IL-10, and (G) IL-33 in cochlear lysate from FKN WT mice subjected to either no-noise exposed (NNE) or noise-exposed and TT injected with vehicle at 1 day post exposure (NE; vehicle) or noise-exposed and TT injected with sFKN peptide at 1 day post exposure (NE; sFKN). N=3-4 biological samples per experimental group and each biological sample is a pool of five cochleae that is ran in triplicates. Values are means ± SD. *P < 0.05, ****P < 0.001 between NNE and NE; vehicle. There was no significant difference in the means between NNE and NE; sFKN or NE; vehicle and NE; sFKN experimental groups. 1-way ANOVA, Dunnett’s multiple comparisons test.

### sFKN peptide is bioavailable in the cochlea following transtympanic injection

In order to verify that the TT injected sFKN peptide into the middle ear reaches the inner ear, cochlear protein lysate and perilymph were extracted at different time points after sFKN TT injection in FKN KO mice and sFKN level (cochlear bioavailability) was estimated by ELISA (Fig. 7A) and MALDI-TOF-MS (Fig. 7B and 7C), respectively. Compared to the uninjected or control ears, a significant level of sFKN was detected after 1 hour of TT injection in cochlear protein lysate by ELISA (Fig. 7A) and in perilymph by MALDI-TOF-MS (Fig. 7C). Approximately, 1/10 to 1/25 of the total sFKN peptide (250 ng in 5 µl at a concentration of 50 ng/µl) injected transtympanically into the middle ear reaches inside the cochlea. The sFKN level in the cochlea was maintained for 12 hours post injection after which the level started to decline by 24 hours post injection. Additionally, mouse sFKN conjugated to Alexa Fluor 647 (sFKN-647) was TT injected in FKN WT mice to visualize the spatial distribution of sFKN-647 at 3 hours post injection. Data shows that injected sFKN-647 distributes into the cochlear sensory epithelium, spiral limbus, spiral lamina, and spiral ligament in all three turns at 3 hours post injection (Fig. 7D, right panel) when compared to the uninjected ear (Fig. 7D, left panel). Likewise, elevated mean fluorescence intensity of conjugated sFKN-647 in cochlear perilymph was detected at 3 hours after TT injection (Fig. S13). Notably, increased numbers of CD45 immunolabeled cochlear macrophages were seen in the sensory epithelium and osseous spiral lamina at 3 hours after sFKN-647 TT injection (Fig. 7D, bottom right panel) when compared to uninjected ear (Fig. 7D, bottom left panel). Collectively, these data from different complementary approaches confirm that sFKN peptide is bioavailable in the cochlea for up to 24 hours after TT injection and it is localized to the sensory epithelium and spiral lamina where it restores the noise-damaged IHC ribbon synapses and function.

**Fig. 7.**
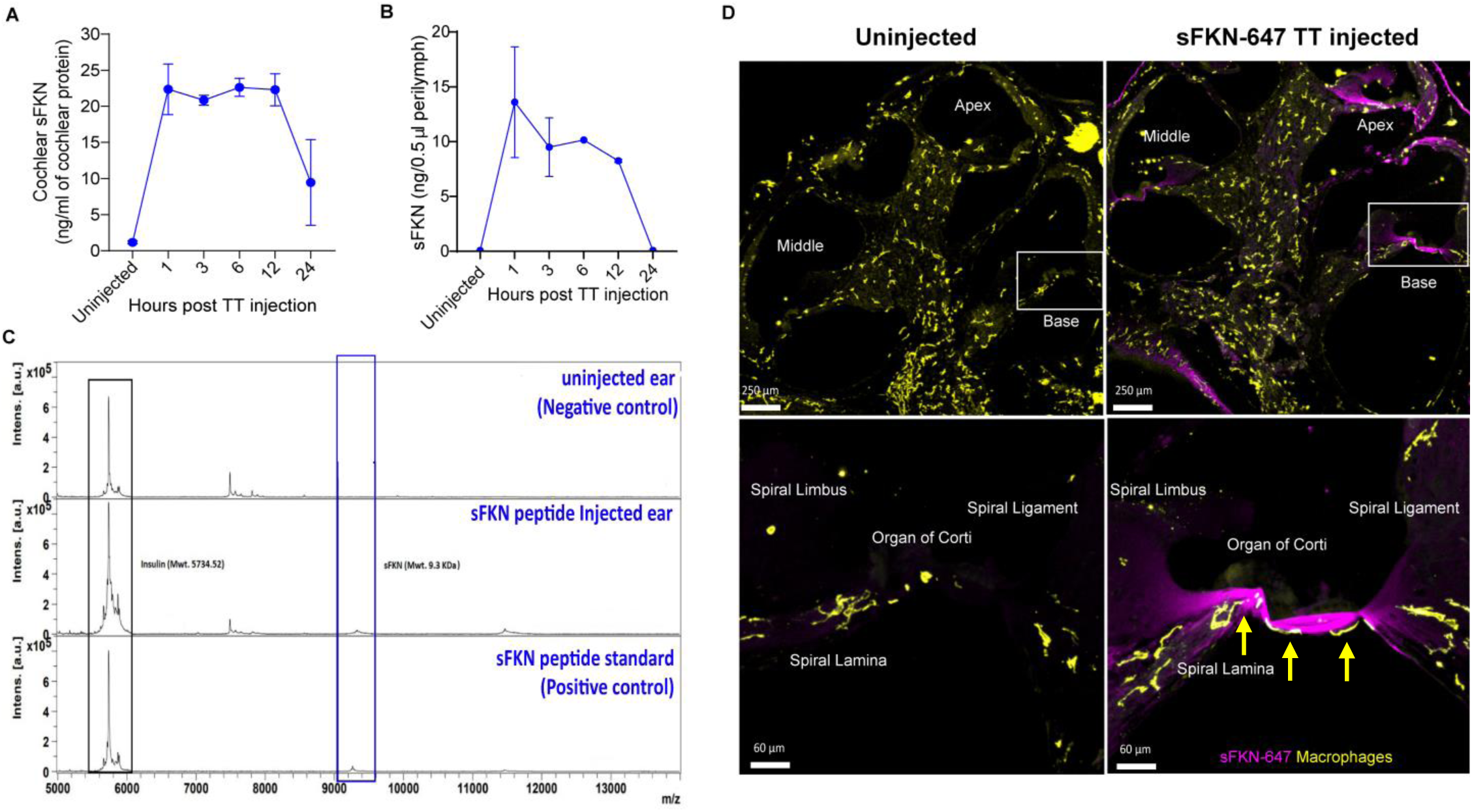
Temporal and spatial bioavailability of sFKN peptide after TT injection in FKN KO and FKN WT mice. Estimation of levels of sFKN peptide in (A) cochlear protein lysate by ELISA and (B) cochlear perilymph by MALDI-TOF-MS at different time points after TT injection in FKN KO mice. N=2 biological replicates per time point (A and B). Values are means ± SD. (C) Representative MALDI-TOF mass spectrometric peaks of sFKN peptide (9.3 kDa) at 3 hours after TT injection in FKN KO mice when compared to uninjected ear (blue rectangle). Peak in the bottom panel represent that of the standard sFKN peptide of 9.3 kDa as positive control. Peaks of the left represent that of insulin peptide as internal control. (D) *Upper panel,* representative images of cochlear midmodiolar crossection at low magnification and *bottom panel,* representative images at higher magnification of the basal cochlear turn (white rectangular box in upper panel) showing the distribution of fluorescent conjugated sFKN-647 (magenta) in the sensory epithelium, spiral limbus, osseous spiral lamina and spiral ligament of uninjected and TT injected FKN WT mice. Notably, more CD45-immunolabeled macrophages (yellow cells, also see yellow arrows) are found in the sensory epithelium at 3 hours after sFKN-647 TT injection (right, bottom panel).

### Working Model

**Fig. 8.**
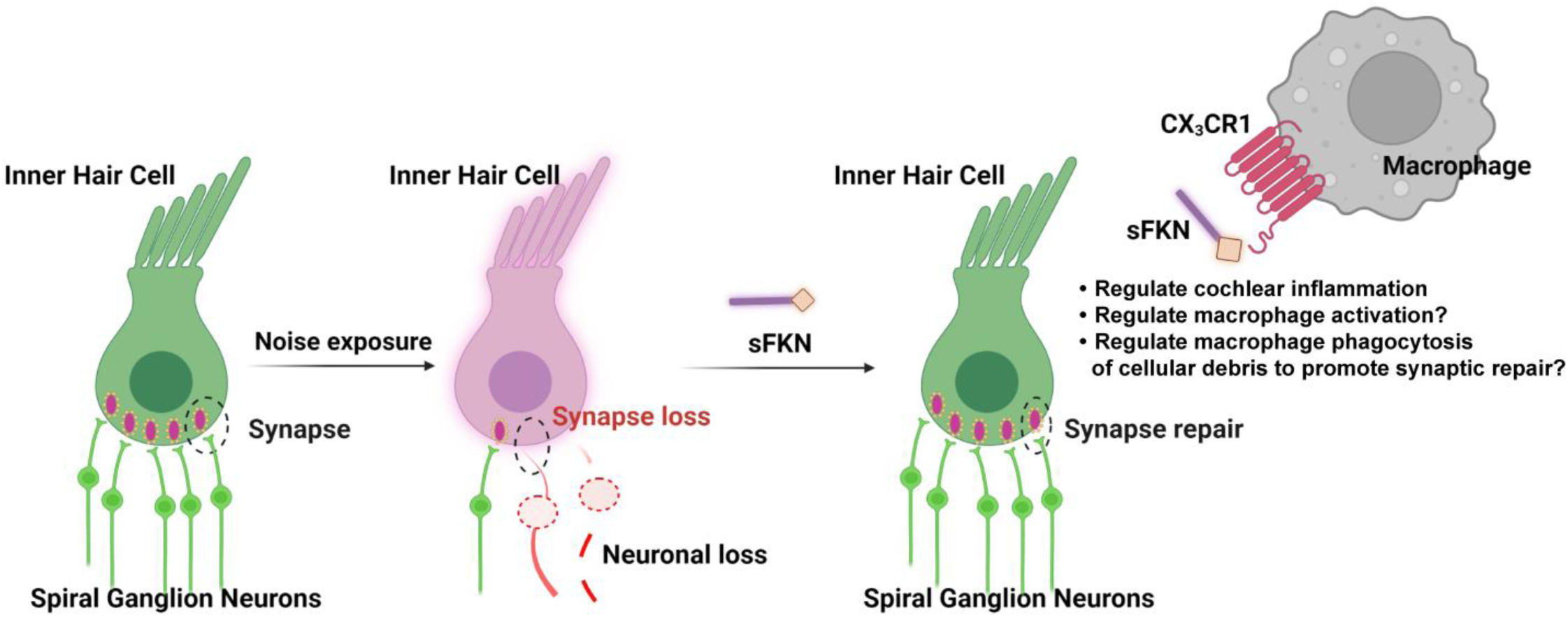
Working model for soluble FKN restores IHC ribbon synapses after NICS. Prolonged or loud exposure to noise results in rapid loss of IHC ribbon synapses known as cochlear synaptopathy. Locally (transtympanically) delivered immune factor, soluble FKN reaches the cochlear sensory epithelium of the inner ear and is effective in restoring the noise-induced loss of IHC ribbon synapses and hearing via cochlear macrophages expressing CX_3_CR1. The precise molecular mechanism by which sFKN-macrophage interaction restore damaged ribbon synapses remains to be elucidated. Figure is created with BioRender.com.

## DISCUSSION

In this study, we specifically assessed the efficacy of FKN isoforms i.e., soluble, and membrane-bound in restoration of loss of IHC ribbon synapses and hearing after synaptopathic noise trauma. The present study offers compelling evidence that soluble FKN peptide is more effective than membrane-bound FKN in restoring lost IHC ribbon synapses and hearing in a mouse model of NICS. IHC synaptic repair with sFKN requires the presence of cochlear resident macrophages and attenuation of inflammation due to NICS without altering the expression of CX_3_CR1 receptor on cochlear macrophages. We also report that sFKN peptide is bioavailable in the cochlea for 24 hours after a single transtympanic injection and it is localized to the sensory epithelium and spiral lamina where it restores the synaptopathic noise-damaged IHC ribbon synapses and function (Fig. 8). While the mechanism by which sFKN restore ribbon synapses remains to be elucidated, our findings suggest that sFKN therapy could be beneficial for patients with NICS, and potentially for other patients with cochlear synaptopathy that affect understanding of speech or central auditory processing disorders.

FKN is a 373-aa protein with a single transmembrane domain that can undergo proteolytic cleavage to release sFKN into the extracellular environment (*20*). In the CNS, both mFKN and sFKN are primarily produced by neurons, and by binding CX_3_CR1 on microglia, they are thought to regulate key aspects of microglial physiology (*56, 77*). One of the main tasks of FKN in neuron–microglia interactions is to suppress the activation of microglia (*72, 78*). Supporting this notion, exogenous delivery of sFKN has been shown to decrease microglia activation as well as neurological deficits in animal models of Parkinson’s disease, stroke, retinitis pigmentosa, and diabetic retinopathy (*39, 44, 47, 50, 52*). In the cochlea, FKN is constitutively expressed by sensory SGNs, IHCs and certain supporting cells (*16*) and sFKN is released after selective cochlear hair cell ablation (*16*), or after NICS (Fig. S8) or it is upregulated after a PTS-imparting noise trauma (*79*). Moreover, in the absence of CX_3_CR1 on cochlear macrophages there is an impairment in spontaneous synaptic repair and enhanced loss of SGNs and IHCs after NICS (*12*). Based on these data, we boosted the levels of FKN in the cochlea at 1-day after a synaptopathic noise trauma for 2 hours at 93 dB SPL with the hope that it would restore the damaged IHC ribbon synapses and hearing function. Single transtympanic injection of sFKN indeed restored noise-damaged synapses and hearing function and it did so in the presence of cochlear macrophages and by attenuating inflammation due to NICS. Remarkably, synaptic repair was seen with the treatment of sFKN, but not with full-length membrane-bound FKN (mFKN) which strongly corroborates the pre-clinical animal studies in the CNS (*45, 52*). This could be due to differences in the bioavailability of the transtympanically injected peptides where sFKN as being smaller peptide (9.3 kDa) than mFKN peptide (34 kDa) crosses the round and/or oval window membrane and is better able to reach cochlear resident macrophages. Of note, we found using three complimentary approaches i.e., ELISA, MALID-ToF-MS and conjugated sFKN-647 that TT injected sFKN peptide reaches the cochlea and it is bioavailable for 24 hours post injection that might be sufficient to ligate to CX_3_CR1 on cochlear resident macrophages to activate downstream signaling in those macrophages and restore synapses. Alternatively, it could be that ligation with only sFKN but not the mFKN allows for CX_3_CR1 internalization to the cytoplasm following ligation to affect downstream signaling pathway (*45, 80, 81*). Although ligation of mFKN with CX_3_CR1 is possible through the chemokine domain but its permanent attachment to the membrane may prevent subsequent internalization to the cytoplasm of macrophages, thereby altering any downstream effectors. Therefore, only the soluble domain of FKN can be readily internalized with CX_3_CR1 ligation. This notion that CX_3_CR1 actively removes sFKN from the surrounding environment is supported by Cardona et al., 2008, where the authors reported substantially more circulating or soluble FKN in CX_3_CR1 ^-/-^ mice (*82*). By using FKN KO mice, we were able to isolate the effects of individual FKN isoform in synaptic repair in response to NICS. sFKN restored noise-damaged ribbon synapses and hearing function in both FKN KO and WT mice. While synaptic repair was near to complete in the FKN KO mice, it was incomplete yet significant in the FKN WT mice. This effect could be due to the pharmacological ceiling effect of exogenous sFKN in the wildtype mice which also releases endogenous sFKN after NICS (Fig. S8) (*44*) which suggest that sFKN pharmacological potency or dose response studies are further warranted.

Activated macrophages are associated with damaged ribbon synapses in response to NICS, given their immediate migration into the IHC synaptic region, upregulation of inflammatory genes, and phagocytosis of debris (*12, 66, 68*). These macrophage activities are beneficial in NICS by promoting IHC synaptic repair, as pharmacological ablation of macrophages has been demonstrated to impair spontaneous synaptic repair (*13*). Here, we found evidence of macrophage activation during NICS, as illustrated by the presence of macrophages in the sensory epithelium near to the IHC synaptic region, consistent with our previous report (*12*). Interestingly, these macrophages were absent or reduced in numbers in the cochlear sensory epithelium of mice lacking FKN, suggesting a critical role of endogenous FKN in localizing macrophages to the sensory epithelium in response to NICS. Likewise, more macrophages were localized to the sensory epithelium with the overexpression of conjugated sFKN-647 peptide in the undamaged cochlea, further supporting the notion that FKN may be the potential chemoattractant for macrophages at least during acute cochlear injury such as NICS. Importantly, FKN contributes to wound healing or repair by recruiting macrophages to the site of injury in a mouse model of skin excision (*83*). Notably, depletion of cochlear resident macrophages abrogated sFKN-mediated restoration of IHC ribbon synapses and function. This indicate that sFKN restores synapses after NICS in a macrophage-dependent manner. We have previously reported that CX_3_CR1, specifically expressed by macrophages is necessary for spontaneous repair of IHC ribbon synapses after NICS as deficiency of CX_3_CR1 has been demonstrated to impair synaptic repair (*12*). Here, we found that overexpression of sFKN did not alter the mRNA expression of cochlear CX_3_CR1. Consistent with this, it has been reported that sFKN therapy also does not affect the protein levels of CX_3_CR1 in the brain (*45*). While sFKN does not alter the levels of CX_3_CR1 receptor, whether CX_3_CR1 activation is critical for sFKN-mediated synaptic repair remains to be elucidated. Studies in the CNS have demonstrated an upregulation of lysosomal pathways in microglia with sFKN therapy, a prominent feature of activated microglia, implicating that these cells may more efficiently digest or phagocytose cellular debris thus favoring preservation of neurons (*52*). Moreover, sFKN therapy attenuates the induction of proinflammatory cytokines in microglia, in other words it represses microglial response, as prolonged or uncontrolled induction can result in chronic inflammation, thereby leading to neurodegeneration (*42, 45, 84, 85*). Here, we also found evidence for macrophage activation during NICS, as demonstrated by upregulation of IFN ß, IL-6, IL-2, 1L-23 cytokines in whole cochlear protein lysates. Remarkably, overexpression of sFKN attenuated or maintained the production of these cytokines to a similar level found in the undamaged cochlea. Our data are consistent with the CNS studies and indicate a plausible mechanism by which sFKN restore synapses partly by modulating cochlear inflammation, although how does inflammation disrupts intact synapses or impairs synaptic repair remains to be determined. While our data indicate that macrophages are obligatory for sFKN to mediate synaptic repair, the effect of sFKN on phagocytic or lysosomal activation of cochlear macrophages and the mechanisms by which sFKN-macrophage interaction promote synaptic repair remains to be elucidated.

Our data provide convincing evidence for the presence of sFKN peptide in the cochlea after a single TT injection in the middle ear. Targeted or local delivery like TT injection is preferable to systemic administration because this can circumvent the side effects and other undesirable effects of the FKN i.e., macrophage activation in peripheral organs or in circulation (*86*), which in turn may allow for higher local doses and longer treatment durations. Actually, repeated TT injections of gentamicin antibiotics or steroids are routinely done in clinical practice for treating Meniere’s disease or idiopathic sudden sensorineural hearing loss, respectively (*87*). TT injection is also preferred over invasive round window or semicircular canal delivery that leads to inflammation and immune response thereby, requiring further treatment with steroids. Notably, transtympanically injected sFKN peptide was distributed throughout all the cochlear turns and was specifically localized to the sensory epithelium and spiral lamina in each turn where the IHC ribbon synapses are damaged after NICS. This is particularly interesting because sFKN has direct access to the damaged site to promote synaptic repair and was also associated with increased macrophages in the same cochlear region. Both ELISA and mass spectrometry data revealed the cochlear bioavailability of sFKN peptide with sustained levels for up to 12 hours after which levels started to reduce. This reduction in sFKN levels could be due to proteolytic degradation of the peptide by proteases and peptidases present in the inner ear fluids (*88*). This could be a limitation of FKN peptide-based therapy however, this critical data has opened new avenues for the development of better analogues of sFKN peptide resistant to proteolytic degradation or small molecule agonists of CX_3_CR1 (FKN-mimetic) with desirable bioavailability. Gene therapy for sFKN can also be employed to achieve prolonged bioavailability of sFKN (*45, 47, 52, 55, 84, 85*) but, cochlear delivery and cell specific transduction of genes is still challenging and entails a risk for immunosuppression. On the contrary, as peptides are short acting, their therapeutic efficacy, potency, and toxicity can be tightly regulated versus gene therapy. In addition, modulation of FKN signaling with peptides could provide a new prevention and treatment platform for noise-induced hearing loss since peptides present an opportunity for therapeutic intervention that closely mimics natural pathways. Nonetheless, we consider that reported bioavailability of sFKN in the present study is sufficient to ligate CX_3_CR1 and recruit macrophages to the damaged IHC synaptic region in the sensory epithelium to activate downstream signaling pathways like phagocytosis and resolution of inflammation that promotes restoration of noise-damaged ribbon synapses and hearing function.

Currently, there are no FDA-approved drugs or treatments for NICS that elicit regeneration of lost auditory nerves and replenish their synaptic connections with surviving hair cells and restore hearing. Therefore, there is a great need to identify novel molecules to develop therapeutic targets to restore synapses and hearing in NICS. Our findings have identified a novel immune pathway, fractalkine signaling axis and they provide a robust proof-of-principle that sFKN can be developed as an immunotherapy that would be beneficial for patients with NICS, and potentially for other patients with cochlear synaptopathy that affect understanding of speech or central auditory processing disorders. Clinically, sFKN could be administered either alone or together with potential neurotrophins like NT-3 or BDNF which partially restore synapses in animal models of NICS (*7–11*). Due to the anti-inflammatory and regulating macrophage activation properties, sFKN can also be tested against other forms of sensorineural hearing loss such as age-related hearing loss and cochlear synaptopathy that are associated with inflammation, vascular damage, and macrophage dysregulation (*89*).

### Limitations of the study

The present study did not examine the pharmacological potency (dose response), dosage, therapeutic window, and toxicology of soluble FKN in NICS. Additionally, the effect of sFKN in modulating macrophage density, morphological activation (transformative index), phagocytic index and inflammation in NICS was not undertaken in the present study. Before sFKN can be used clinically for noise-induced cochlear synaptopathy and hidden hearing loss these studies are warranted.

## MATERIALS AND METHODS

### Mice

Study used in-house bred age-matched FKN wild-type mice (hereafter denoted as FKN WT) (Jackson Laboratories, Bar Harbor, Maine, stock number, 000664) and CX_3_CL1^-/-^ or FKN knockout mice (hereafter denoted as FKN KO) (*90*), gifted by Dr. Astrid Cardona, University of Texas, Department of molecular Microbiology and Immunology, College of sciences, San Antonio, Texas) on a C57BL/6J background. Efforts were made to minimize animal suffering and to reduce the numbers of animals used. Animals were housed under a 12-hour light/12-hour dark cycle and fed ad libitum, unless specified. All aspects of animal care, procedure and treatment were carried out according to guidelines of the Animal Studies Committee of Creighton University, Omaha, NE and Rutgers University, Piscataway, NJ.

### Chemicals and peptides

Chemicals and peptides were purchased from Fisher Scientific, Sigma-Aldrich or R&D systems until mentioned otherwise. Poloxamer 407 (Sigma-Aldrich, cat. #16758), recombinant control peptide (R&D systems, cat. #4460, Accession # P01863) is mouse melanoma cell line, NS0-derived mouse IgG2A protein of 26.7 kDa and 232 amino acids, recombinant full length membrane-bound Fractalkine (R&D systems, cat. #472-FF, Accession # O35188) is sf21 (baculoviris)-derived mouse CX3CL1/Fractalkine protein of 34 kDa and 312 amino acids and recombinant soluble Fractalkine (R&D systems, cat. #571-MF, Accession# AAB71763) is *E. coli*-derived mouse CX3CL1/Fractalkine peptide from amino acid 25 to 105 of 9.3 kDa and 80 amino acids.

### Transtympanic (TT) Injection

Lyophilized powders of the peptides indicated above were reconstituted with sterile 1xPBS, pH 7.4 (Fisher Scientific, cat. #BP661-10) to an equimolar stock concentration, aliquoted and stored at −80°C until needed for experiment. Immediately before the TT injection, aliquots were mixed with an equal volume of sterile 18% poloxamer 407 hydrogel and placed on ice to maintain the fluidity and to prevent gel formation. The poloxamer hydrogel is liquid at room temperature or below, allowing TT injection, but transitions to a gel at body temperature, providing a prolonged residence time in the middle ear (*91*). Mice were anesthetized with an intraperitoneal injection of Ketamine/Xylazine (Patterson Veterinary) cocktail (100 and 20 mg/kg, respectively; dose: 0.1 ml/20 gm body weight), following procedures described in our previous publications (*12*). Anaesthetized mouse’s ear was brought under the focus of a stereo dissection microscope (Vermont Optechs, Inc., Maine, USA). A speculum was placed in the outer ear. Carefully an exhaust hole was made in pars tensa of the tympanic membrane using 1 ml insulin syringe needle (Excel, cat. #26029). The injection hole of ∼ 0.5mm was made ∼90° angle opposite to the exhaust hole below the pars flaccida. Vehicle or peptide mixed with vehicle was injected by inserting the needle (31 gauge, Hamilton, cat. #7762-04) attached to a 10 µl Hamilton syringe (Hamilton, cat #) into the injection hole and the content were slowly plunged near the round window. The mouse was placed on a heating pad with the injected ear up until it recovered from the anesthesia.

### Study design

FKN WT and KO mice of both sexes at 5-6 weeks of age were used for study. To avoid the saturation effect of the cochlear endogenous FKN ligand, FKN KO mice were used to test the efficacy of FKN isoforms in restoration of ribbon synapses after NICS. Following assessment of baseline hearing sensitivity at 1-day pre noise exposure (PNE), mice were exposed to a synaptopathic noise level of 93 decibel sound pressure level (dB SPL) at 8-16 kilohertz (kHz) for 2 hours (*13*). Degree of hearing loss was measured at 1-day post noise exposure (DPNE), immediately after which the noise exposed FKN WT mice were TT injected with the sterile vehicle (18% Poloxamer 407 hydrogel). Noise exposed FKN KO mice were injected either with vehicle (18% Poloxamer 407 hydrogel), control peptide, membrane-bound FKN peptide (mFKN) or soluble FKN peptide (sFKN). All peptides were TT injected at a concentration of 50 ng/µl based on the reported EC_50_ in CNS studies (*43–53*) in a total injection volume of 5 µl (total peptide delivered = 250 ng) and all TT injections were performed once and unilaterally. Degree of recovery of hearing thresholds shifts was measured at 14 −15 DPNE, following which the mice were euthanized and isolated cochleae were microdissected and processed for various assays indicated below. Age-matched unexposed and uninjected FKN WT and FKN KO mice served as controls (Fig.1A).

### Noise exposure

Awake, unrestrained mice were exposed for 2 hours to an octave band-noise (8-16 kHz) at graded noise levels (93 dB SPL). Noise exposures were performed in a foam-lined, single-walled soundproof room (WhisperRoom Inc.,TN). Briefly, mice were placed singly or in pairs in modified subdivided cages (food, water, bedding removed) positioned up to two cages at once directly under an exponential horn (JBL). Noise was generated digitally using custom Labview routines and a Lynx E22 sound card (running on a PC), filtered pure tone (8–16 kHz), and amplified (D-150A power amplifier, Crown Audio) that drive the speaker through an exponential horn (JBL). Prior to each exposure, the noise was calibrated to target SPL using a quarter-inch condenser microphone (PCB). Sound pressure levels varied by ±1 dB across the subdivisions in the cage. Unexposed mice served as age-matched sham controls.

### Electrophysiological recordings

Auditory brainstem responses (ABR) and distortion product otoacoustic emissions (DPOAE) were measured by an observer who was “blinded” to the experimental conditions of each animal. Mice were anesthetized with an intraperitoneal injection of Ketamine/Xylazine (Patterson Veterinary) cocktail (100 and 20 mg/kg, respectively; dose: 0.1 ml/20 gm body weight), following procedures described in our previous publications (*12*). ABRs thresholds and DPOAEs levels were measured prior to noise exposure, at 1 day after noise exposure (to verify degree of hearing loss) and at 2 weeks after noise exposure, to quantify recovery of hearing.

#### ABRs

Anesthetized mice were placed on a heating pad set at 37°C and eyes were lubricated with an ophthalmic ointment (Artificial Tears) to avoid drying due to anesthesia. Subcutaneous needle electrodes were placed behind the right pinna (reference) and vertex (active). A ground electrode was placed near the tail of the mouse. A TDT Multi Field (MF1) speaker was set at 10 cm from the right pinna and calibrated with PCB ¼” free field calibration microphone before recordings. Stimuli of 5-ms tone pips (0.5 ms cos^2^ rise-fall) were delivered at 21/s with alternating stimulus polarity using an RZ6-A-P1 Bioacoustic system and Medusa4Z pre-amplifier (Tucker-Davis Technologies, Alachua, FL). Recorded electrical responses at a sampling rate of 12 kHz were filtered (300 Hz to 3 kHz) and averaged using BioSigRZ software (Tucker-Davis Technologies, Alachua, FL). The sound level was decreased in 5-dB steps from 100 dB Sound Pressure Level (SPL) down to 10 dB SPL. At each sound level, 512-1024 responses were averaged, and response waveforms are discarded as artifacts if the peak-to-peak voltage exceeded 15 μV. Thresholds at 8.0, 11.2, 16.0, 22.6, and 32.0 kHz were determined by a single observer who noted the lowest sound level at which a recognizable Peak I (PI) waveform could be obtained. Waveforms were confirmed as auditory-evoked responses by their increasing latency and decreasing amplitude as the intensity of the stimulus is lowered. ABR thresholds refer to the lowest SPL that can generate identifiable electrical response waves. These threshold values (actual or assigned) were used to calculate the mean ABR thresholds at each stimulus frequency. Threshold shifts were calculated by subtracting baseline thresholds from thresholds at 1 day and 15 days post noise exposure at all the stimulus frequencies.

#### Input/output (I/O) neural response

Methods followed those described in Kaur et al., 2019 (*12*). Briefly, ABR Peak I component was identified and the peak to trough amplitudes and latencies were computed by off-line analysis of stored ABR waveforms. ABR Peak I amplitude and latency versus-stimulus level (ABR I/O) data were computed at 8, 16, 22.6 and 32 kHz for sound levels ranging from 100 dB SPL up to the hearing thresholds, unless otherwise specified.

#### DPOAEs

Anesthetized mice were placed on a heating pad set at 37°C and eyes were lubricated with an ophthalmic ointment (Artificial Tears) to avoid drying due to anesthesia. Stimuli were presented at 5 –40 kHz and delivered to the right ear by a custom coupling insert RZ6-A-P1 bioacoustic system (Tucker-Davis Technologies, Alachua, FL). Electrostatic loudspeakers (EC1) were calibrated with ER10B+ Etymotic low noise probe and microphone before recordings. Distortion product (DP) grams (2f1-f2) were obtained for f2 ranging from 5 to 40 kHz, with a frequency ratio of f2/f1 of 1.2 and L1–L2 = 10 dB. Recordings were performed using BioSigRZ software (Tucker-Davis Technologies, Alachua, FL). Following ABR/DPOAE recordings, animals were placed on a heating pad at 37°C and monitored until they regained activity and then returned to animal research facility.

### PLX5622-induced depletion of cochlear macrophages

To determine whether macrophages are necessary for sFKN-mediated synaptic repair after NICS, FKN WT mice were fed on either standard nutritionally-complete rodent diet (AIN-76A formulation, referred to as control chow) or on AIN-76A chow that contained 1.2 gm/kg of PLX5622 (hemifumarate salt, 99.30% purity, cat. # HY-114153, MedChemExpress, NJ, referred to as PLX5622 chow). PLX5622 is a selective antagonist of Colony Stimulating Factor 1 Receptor (CSF1R), that regulates macrophage development, survival, and maintenance (*92*). Both control and PLX5622 chows were formulated, irradiated, and supplied by Research Diets, Inc., New Brunswick, NJ. Following 2 weeks on respective chows which led to a robust elimination of cochlear resident macrophages, both control and PLX5622 fed FKN WT mice were exposed to noise after recording the baseline hearing sensitivity followed by the TT injection of either vehicle or sFKN peptide at 1 DPNE, and the special chows were maintained until the end of the experiment for sustained macrophage depletion. Hearing function was assessed again at 15 DPNE followed by euthanasia, and processing of isolated cochleae for synaptic immunolabeling. Feeding mice with PLX5622 chow for 15 days does not affect the normal hearing sensitivity, as shown in our previous work (*13*).

### Tissue harvesting and processing

Mice were deeply anesthetized with lethal doses of pentobarbital sodium (Trade Name-Fatal Plus, Patterson Veterinary). Prior to respiratory arrest, mice were perfused by transcardiac route with phosphate buffered saline (PBS) (Fisher Scientific, cat. # BP661-10) or 4% paraformaldehyde (Fisher Scientific, cat. # 50980495) in 0.1 M phosphate buffered solution and temporal bones were harvested. For fluorescent immunolabeling of cochlear synapses, excised temporal bones were post fixed in 4% paraformaldehyde for 20 minutes on ice and decalcified in 0.1 M ethylenediaminetetraacetic acid (EDTA, Fisher Scientific, cat. # AC327205-000) overnight for ∼16 hours. For macrophage immunolabeling in cochlear mid-modiolar cryosections, temporal bones were fixed for 3-4 hours at room temperature and decalcified for 3-5 days. The temporal bones were rinsed thrice with PBS and appropriate fluorescent immunohistochemistry was performed.

### Immunohistofluorescence

Microdissected cochlear whole mounts or frozen mid-modiolar cross sections (20 μm) were rinsed with PBS (Fisher scientific, cat. # BP661-10) at least 3 times and incubated at room temperature for 2 h in blocking solution containing 5% normal horse serum (Sigma-Aldrich, Cat. # H0146) in 0.2% Triton X-100 (Sigma-Aldrich, cat. # A16046AP) in PBS. For synaptic immunolabeling, cochlear microdissected whole mounts were incubated overnight at room temperature with combinations of the following primary antibodies: CtBP2 mouse (BD Biosciences, cat. # 612044; RRID:AB_399431; 1:200), GluA2 mouse (EMD Millipore, cat. # AB1506; RRID:AB_11212990; 1:100), Myosin 7a rabbit (Proteus Biosciences, cat. # 25-6790; RRID:AB_2314838; 1:500). For macrophage and neuron immunolabeling, cochlear frozen mid-modiolar cross sections or whole mounts were incubated overnight at room temperature with combinations of following primary antibodies: goat anti-CD45 (R&D Systems, catalog # AF114; RRID:AB_442146; 1:100), mouse anti-Neurofilament 165 (NF-H) (Developmental Studies Hybridoma Bank, cat. # 2H3C, 1:100) and mouse anti-Tuj1 (Covance, cat. #MMS-435P, 1:500). Following incubation with primary antibodies, specimens were rinsed 5 times in PBS and treated for 2 h at room temperature in species-specific secondary antibodies conjugated to either DyLight 405 (1:500, Jackson ImmunoResearch Laboratories Inc.), or AlexaFluor-488, −546, −555, or −647 (1:500; Invitrogen, Life Technologies). Tissue was mounted in glycerol:PBS (9:1) and coverslipped before confocal imaging. Tissue samples were batch processed using the same reagent solutions for all experimental groups.

### Confocal Imaging

Three- or four-color fluorescence imaging was performed using a Zeiss LSM 700 laser scanning confocal microscope (Carl Zeiss Microscopy 700, Jena, Germany). Z-series images using 5x,10x, 20x, 40x or 63x objectives were obtained. Image processing and quantitative analyses were performed using IMARIS (version 9.9.0, Oxford Instruments) and Volocity 3D image (version 6.5.1, PerkinElmer) software.

### Synapse count

Two to three confocal z-stacks were obtained using a high-resolution oil immersion objective (63x with 1.4 numerical aperture) and a digital zoom of 1.5 from apical, middle, and basal regions per cochlear whole mount. Each z-stack spanned the entire synaptic pole of 8-10 IHCs in the z-dimensions, with z-step-size of 0.3 μm, from the apical portion of the IHC to nerve terminal in the habenula perforata region. IMARIS was used for 3D analysis of individual and juxtaposed/paired pre- and postsynaptic counts. Thresholds for all the channels were adjusted in such a way the background is reduced without losing any prominent fluorescence and two close fluorescent punctae are not merged. Total CtBP2 punctae, total GluA2 punctae and paired CtBP2 and GluA2 fluorescence surface were counted per image. Total counts were divided by the number of surviving IHC to report CtBP2 punctae per IHC, GluA2 punctae per IHC and paired ribbon synapses per IHC.

### Macrophage count

To assess macrophages per 100 µm of sensory epithelium, CD45-labeled macrophages were counted from 40X objective images taken from the apex, middle and basal region of cochlear whole mounts (*16*).

### ELISA

Protein extracted from cochlear lysate was estimated using BCA kit (Thermo Scientific, cat. # 23225). Level of cochlear endogenous FKN (R&D systems, #DY472) or TT injected (exogenous) sFKN peptide (Invitrogen, cat. #EMCX3CL1) was estimated from 50 µg of cochlear protein lysate by following the manufacture’s protocol. Briefly, cochlear protein lysates (samples) were loaded along with the sFKN standard in a 96 well plate coated with capture antibody. Plate was incubated for 150 min at room temperature. After washing, the wells were equally treated with binding antibody, followed by biotin conjugate, streptavidin-HRP and TMB substrate. The yellow color developed at the end point was read at 450 nm using Varioskan LUX multimode plate reader (Thermo Fisher scientific, #VLBL0TD2). Concentration of sFKN in cochlear protein lysate was determined using the linear equation derived from the standard curve (Fig. S12A).

### Matrix Assisted Laser Desorption Ionization-Time of Flight-Mass Spectrometry (MALDI-TOF-MS)

Cochlear perilymph collected (∼0.5 µL) at different time points after TT injection of sFKN in FKN KO mice were diluted with 4.5 µL of mass spectrometry (MS) grade water (with 0.1% trifluoroacetic acid) to prepare final volume of 5 µL. 1 µL of each diluted perilymph sample were spotted on a MALDI stainless steel plate and 1 µL insulin as internal standard (9 ng/ µL) along with 1 µL of DHB matrix solution (40 mg/mL in acetonitrile: water 1:1 (0.1% TFA) were added on the same spot for each time point. For standard curve, sFKN peptide was dissolve in MS grade water (with 0.1% TFA) at 250 ng/ µL and diluted sub-sequentially two times up to 0.48 ng/ µL, spotted on the MALDI-plate in triplicate and then 1 µL insulin as internal standard (9 ng/ µL) and 1 µL of DHB matrix solution were added on each spot (Fig. S12B).

The samples were dried in room temperature. MALDI-Tof-mass spectroscopy experiments were performed in an Autoflex III MALDI-TOF/TOF mass spectrometer (Bruker Daltonics, Leipzig, Germany) equipped with a 200-Hz Smart beam laser and using the Flex control v.3.4 software. Mass data was collected with manual laser positioning. For each sample 10 spots were chosen, and 1000 laser shots were collected to obtained 10 spectra in linear positive mode. The IS1 voltage was 20 kV, the IS2 voltage was maintained at 18.97 kV, the lens voltage was 5.67 kV, and laser power was 95%. Sample rate and digitizer setting was set to 1.25 and detector gain was 10x. Mass data were collected in the range 5000-14000 m/z. Mass accuracy was calibrated internally with spiked insulin, intensity of internal standard and sFKN was determined from each sample. Standard curve was prepared by plotting Intensity ratio of sFKN/ Insulin in Y axis and concentration of peptide in X-axis and fitting the data with linear equation (Fig. S12B). Mass of peptide in perilymph was then calculated from standard curve equation.

### Fluorescent conjugated sFKN peptide detection in the cochlea after TT injection

Mouse sFKN Alexa Fluor-647 (Almac, #CAF-51, 9.9 kDa) was reconstituted in PBS and an equimolar amount to the unconjugated sFKN peptide (i.e., 50 ng/µl) was mixed with sterile Poloxamer 407 hydrogel and was TT injected in FKN WT mice. The cochlea and perilymph were isolated at 3 and 6 hours post injection. Midmodiolar cryosections of cochlea were processed for macrophage immunolabeling as described above and imaged using confocal microscope. The collected perilymph was made up to 50 µl in 1xPBS and the fluorescence intensity or emission was measured at 647 nm using Varioskan LUX multimode plate reader (Thermo Fisher scientific, #VLBL0TD2).

### Luminex assay

Luminex assay was performed to detect the protein levels of various cytokines in cochlear protein lysates using a custom-build multiplex panel as per the manufacture’s protocol (Thermo Fisher Scientific, catalog # PPX-26-MXPRMA9). Briefly, magnetic beads coated with target antibodies were added to the 96-well assay plate and washed thrice with wash buffer. Standards for 26 cytokine and chemokine analytes were mixed and serially diluted to generate standard curves. Cochlear protein lysates at a concentration of 50 µg were used. Antigen standards, blanks, sample lysates were incubated with magnetic beads at room temperature at 500 rpm for 120 minutes in the dark. Then beads were washed thrice to remove the excess lysate/standard. 25 µl of detection antibody was added and incubated in the dark for 30 minutes at room temperature. The beads were washed thrice, 50 µl of streptavidin-PE was added and incubated for 30 minutes in the dark at room temperature. Washed beads were re-suspended in 120 µl of reading buffer and the plate was read using a LuminexTM 100/200TM system (Thermo Fisher Scientific, FLEXMAP 3D). Acquired raw data were analyzed using 5PL algorithm offered by Thermo Fisher Scientific (Procartaplex Analysis App available online). Raw data file from the instrument were fed in the application and the lot numbers of standards were entered. By following the manufacturer’s instruction, the application computes the concentration and displays the graph for individual analytes.

### Statistical analyses

All statistical analyses were performed using Prism GraphPad version 10.0.2. Values are expressed as mean ± standard deviation (SD) across animals in each experimental group unless otherwise stated in the figure legends. Student t-test, or one- or two-way ANOVA was applied as appropriate. Significant main effects or interactions were followed by appropriate post hoc tests. Details on error bars, statistical analysis, numbers of mice, and experimental replicates can be found in results and figure legends. Results were considered statistically significant when probability (P-values) of the appropriate statistical test were less than or equal to the significance level, alpha (α) = 0.05.

## List of Supplementary Materials

### Materials and Methods

#### Real time-quantitative polymerase chain reaction (RT-qPCR)

The total RNA was extracted with Biorad-PureZOL (#7326880). Isolated RNA was treated with DNase 1, RNase-free (Thermo Scientific, #ENO521). The first strand cDNA was synthesized from 1 µg of total RNA using high-capacity RNA-to-cDNA kit (Applied Biosystems, #4388950). About 100ng/2µl of cDNA was used as a template for q-RT-PCR performed using TaqMan^TM^ Fast Advanced Master Mix (Applied Biosystems, #4444557) in ABI PRISM 7500 Fast. Expression level was analyzed using TaqMan® Gene expression assay kit for CX3CR1 and GAPDH (Applied Biosystems, # 4331182). The PCR program was as follows: UNG incubation at 50°C for 2 min, polymerase activation at 95°C for 10 min, Denaturation at 95°C for 15 sec, Anneal and extension at 60°C for 1min, total of 40 cycles were performed. Values were normalized to GAPDH and to the no noise exposed group and represented as fold change.

#### Cochlear hair cell count

Cochlear whole-mounts were processed for immunolabeling for hair cells. Both inner and outer hair cells were identified by their immunoreactivity for Myosin 7A. Hair cells were counted from the apical, middle, and basal region of the cochlea, as recorded in 20X objective images. Data was expressed as inner or outer hair cells in 300 µm of sensory epithelium (*16*).

### Supplementary figures and figure legends

**Fig. S1.**
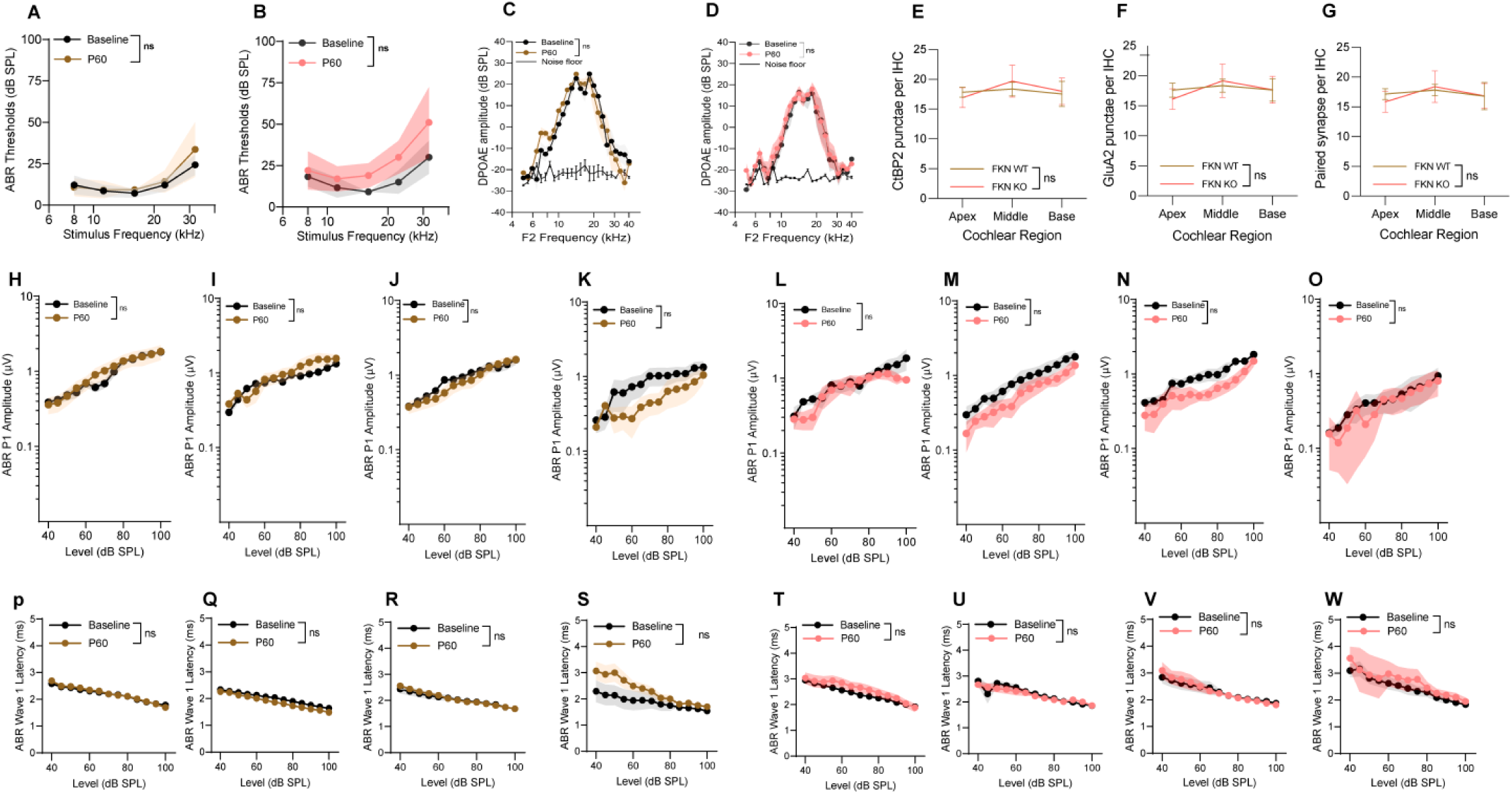
Gross hearing function in unexposed and uninjected FKN WT and FKN KO control mice. ABR thresholds at baseline (P45) and at experimental endpoint (P60) in (A) FKN WT and (B) FKN KO mice. 2-way ANOVA, Sidak’s multiple comparisons. DPOAE levels at baseline (P45) and at experimental endpoint (P60) in (C) FKN WT and (D) FKN KO mice. 2-way ANOVA, Sidak’s multiple comparisons. (E) CtBP2 punctae, (F) GluA2 punctae and (G) paired ribbon synapses per IHC in FKN WT and FKN KO at P60. 2-way ANOVA, Dunnett’s multiple comparisons test. ABR peak I amplitude at baseline (P45) and at experimental endpoint (P60) in FKN WT mice at (H) 8 kHz, (I) 16 kHz, (J) 22.6 kHz and (K) 32 kHz. ABR peak I amplitude at baseline (P45) and at experimental endpoint (P60) in FKN KO mice at (L) 8 kHz, (M) 16 kHz, (N) 22.6 kHz and (O) 32 kHz. ABR peak I latency at baseline (P45) and at experimental endpoint (P60) in FKN WT mice at (P) 8 kHz, (Q) 16 kHz, (R) 22.6 kHz and (S) 32 kHz. ABR peak I latency at baseline (P45) and at experimental endpoint (P60) in FKN KO mice at (T) 8 kHz, (U) 16 kHz, (V) 22.6 kHz and (W) 32 kHz. 1-way ANOVA, Tukey’s multiple comparisons test (H-W). Values are mean ± SD. ns, non-significant comparison between baseline (P45) and at experimental endpoint (P60). N=6-7 mice per genotype.

**Fig. S2.**
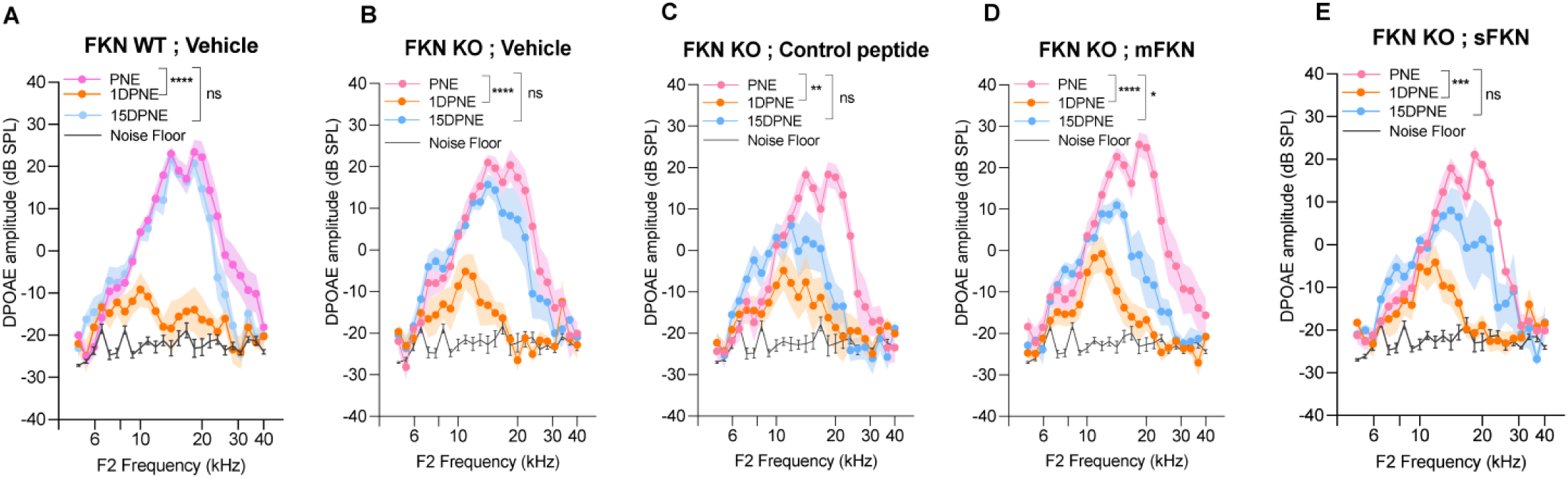
DPOAE amplitude at pre-noise exposure (PNE), 1 DPNE and 15 DPNE. DPOAE levels in (A) FKN WT mice treated with vehicle (N=8), and in FKN KO mice treated with (B) vehicle (N=6) (C) control peptide (N=7) (D) membrane-bound FKN peptide (mFKN) (N=10) and (E) soluble FKN peptide (sFKN) (N=8). Values are means ± SD. *P < 0.05, **P < 0.01, ***P < 0.001, ****P < 0.0001 and ns, non-significant at respective experimental time points. *Represents comparison between PNE and 1DPNE or 15 DPNE as shown with parenthesis, 2-way ANOVA, Dunnett’s multiple comparisons test.

**Fig. S3.**
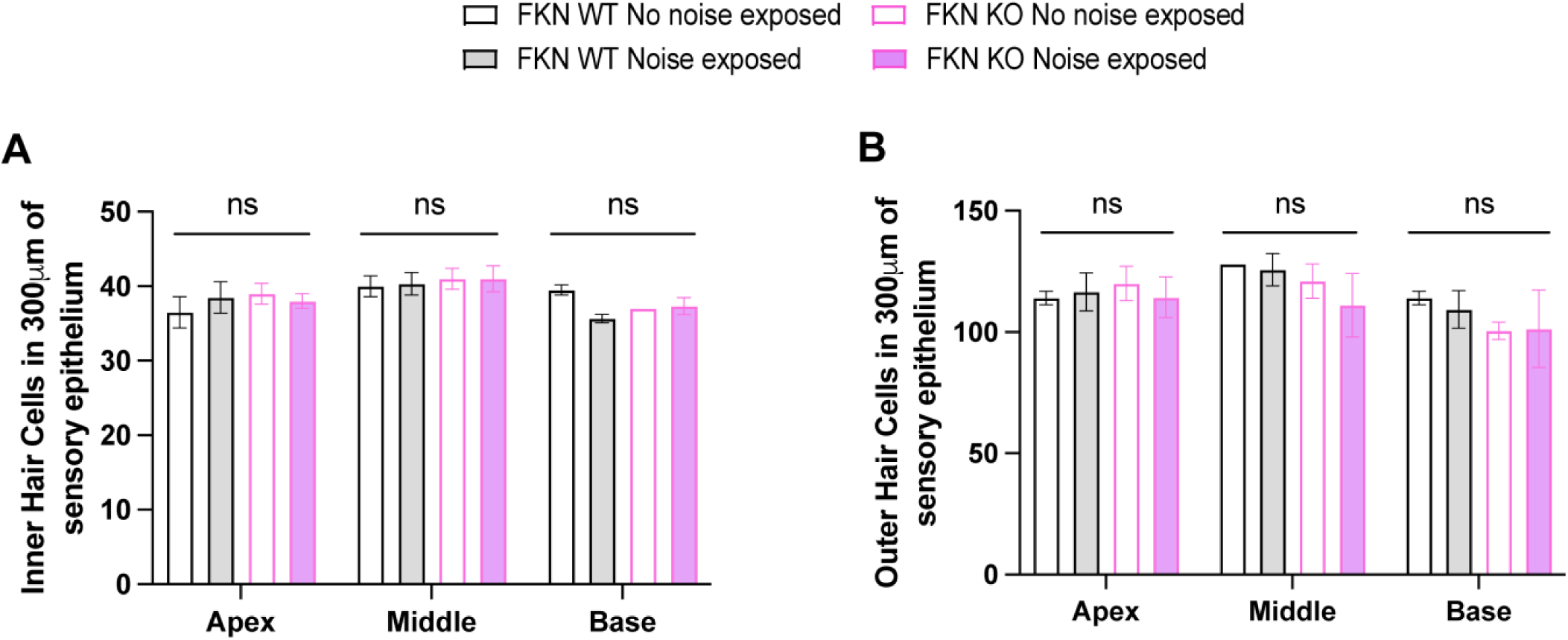
Hair cell density in FKN WT and FKN KO mice after NICS. Density of cochlear (A) inner hair cells and (B) outer hair cells in no noise exposed (open bars) and noise exposed (filled bars) FKN WT and FKN KO mice shows no significant (ns) effect on hair cell survival between the genotypes. N=3 mice per genotype. 2-way ANOVA, Tukey’s multiple comparison test.

**Fig. S4.**
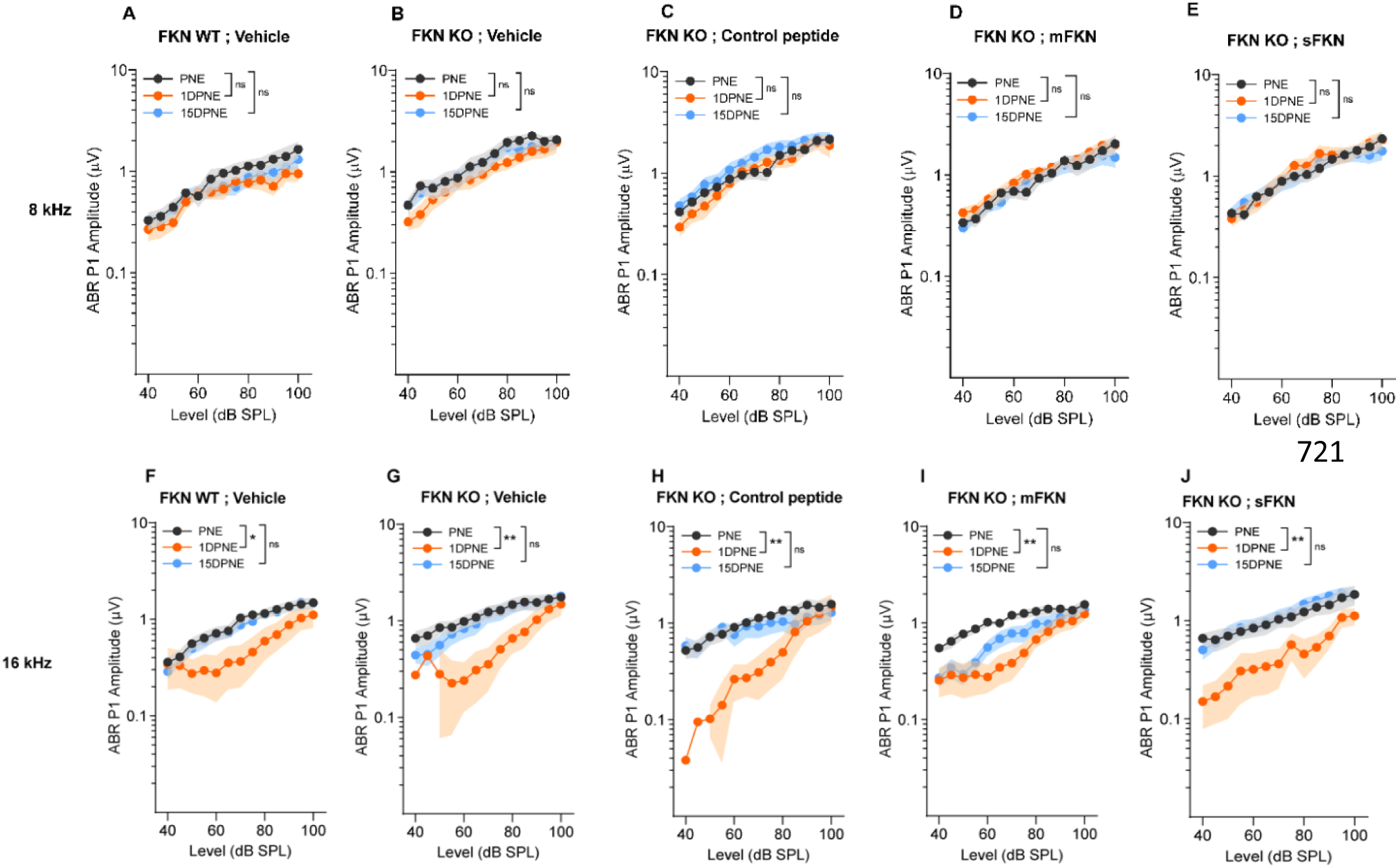
ABR Peak I amplitude at 8 kHz and 16 kHz. ABR Peak I amplitudes at 8 kHz (A) FKN WT mice treated with vehicle (N=8), FKN KO mice treated with (B) vehicle (N=6) (C) control peptide (N=7) (D) membrane-bound FKN peptide (mFKN) (N=10) and (E) soluble FKN peptide (sFKN) (N=8). At 16 kHz (F) FKN WT mice treated with vehicle (N=8), FKN KO mice treated with (G) vehicle (N=6) (H) control peptide (N=7) (I) membrane-bound FKN peptide (mFKN) (N=9) and (J) soluble FKN peptide (sFKN) (N=8). Values are means ± SD. *P < 0.05, **P < 0.01, and ns, non-significant. *Represents the comparison between the experimental time points as indicated with parenthesis. 1-way ANOVA, Tukey’s multiple comparisons test.

**Fig. S5.**
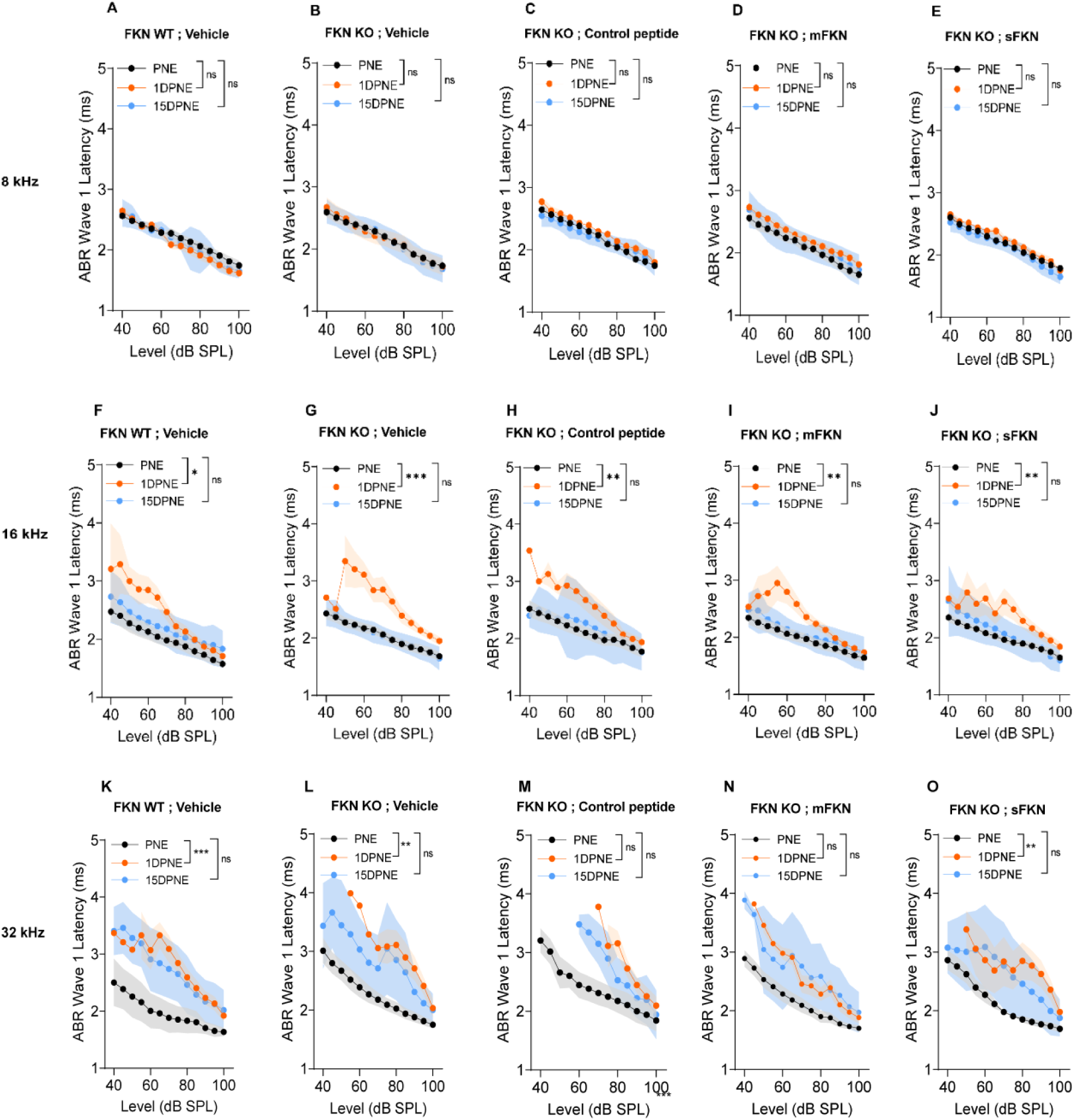
ABR Peak I latency at 8, 16, 32 kHz. ABR Peak I latency at 8 kHz in (A) FKN WT mice treated with vehicle (N=8), and in FKN KO mice treated with (B) vehicle (N=6) (C) control peptide (N=7) (D) membrane-bound FKN peptide (mFKN) (N=10) and (E) soluble FKN peptide (sFKN) (N=8). At 16 kHz in (F) FKN WT mice treated with vehicle (N=8), and in FKN KO mice treated with (G) vehicle (N=6) (H) control peptide (N=7) (I) membrane-bound FKN peptide (mFKN) (N=9) and (J) soluble FKN peptide (sFKN) (N=8). At 32 kHz in (K) FKN WT mice treated with vehicle (N=8), and in FKN KO mice treated with (L) vehicle (N=6) (M) control peptide (N=7) (N) membrane-bound FKN peptide (mFKN) (N=10) and (O) soluble FKN peptide (sFKN) (N=8). Values are means ± SD. *P < 0.05, **P < 0.01, ***P < 0.001, and ns, non-significant. *Represents the comparison between the experimental time points as indicated with parenthesis. 1-way ANOVA, Tukey’s multiple comparisons test.

**Fig. S6.**
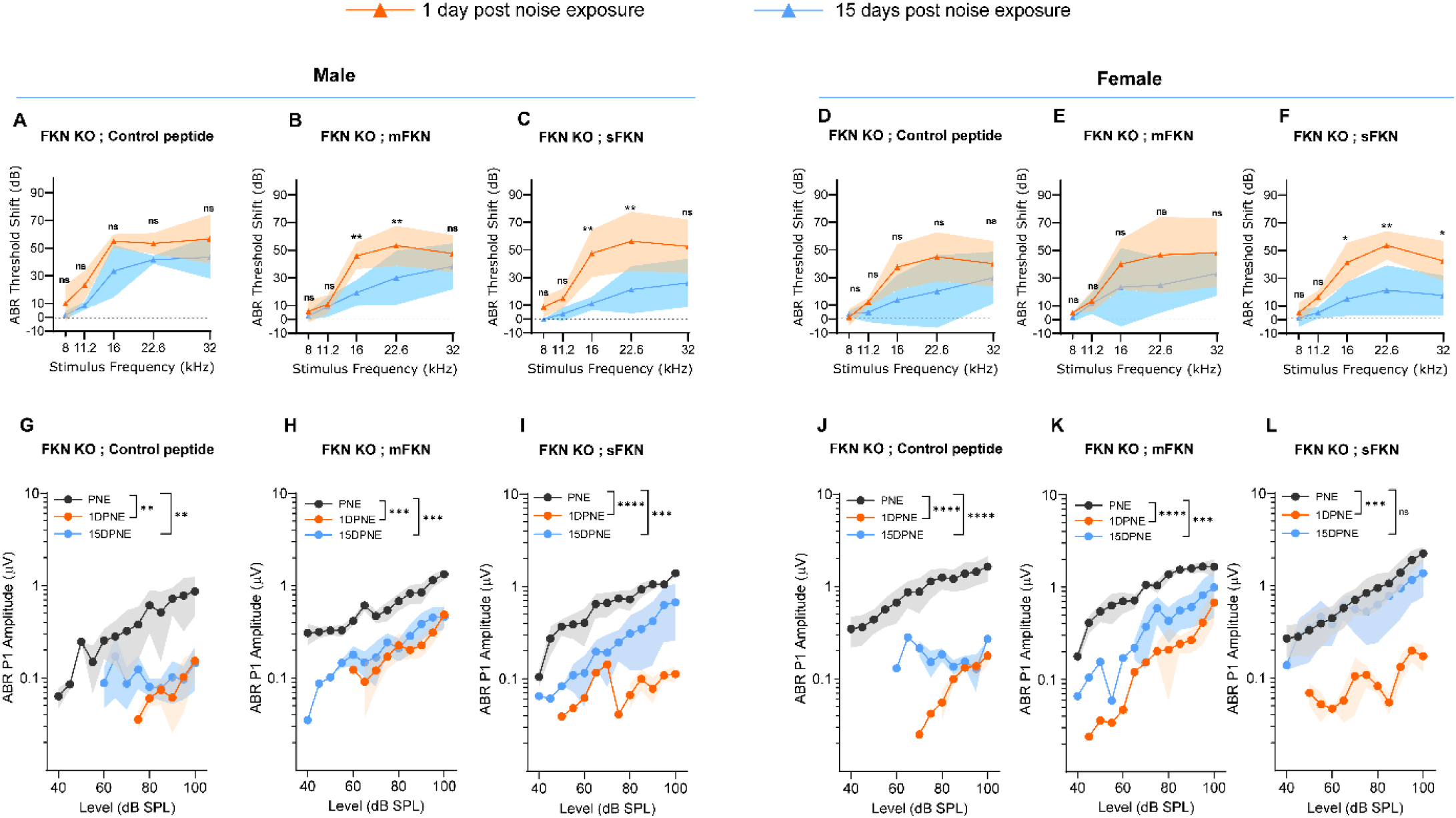
**ABR threshold shifts and Peak I amplitudes in male and female FKN KO mice TT treated with control peptide, mFKN and sFKN peptides**. ABR threshold shifts at 1 DPNE and 15 DPNE at 93 dB SPL from FKN KO male mice treated with (A) control peptide (N=3) (B) membrane-bound FKN peptide (mFKN) (N=5) and (C) soluble FKN peptide (sFKN) (N=4). ABR threshold shifts at 1 DPNE and 15 DPNE at 93 dB SPL from FKN KO female mice treated with (D) control peptide (N=4) (E) membrane-bound FKN peptide (mFKN) (N=3) and (F) soluble FKN peptide (sFKN) (N=4). 2-way ANOVA, Sidak’s multiple comparisons. *Represents comparison between 1 DPNE and 15 DPNE. A-F, Dashed line represent threshold shifts prior to noise exposure (baseline). ABR Peak I amplitudes at 32 kHz from FKN KO male mice treated with (G) control peptide (N=3) (H) membrane-bound FKN peptide (mFKN) (N=5) and (I) soluble FKN peptide (sFKN) (N=4). ABR Peak I amplitudes at 32 kHz from FKN KO female mice treated with (J) control peptide (N=4) (K) membrane bound FKN peptide (mFKN) (N=5) and (L) soluble FKN peptide (sFKN) (N=4). 1-way ANOVA, Tukey’s multiple comparisons test. *Represents the comparison between the experimental time points as indicated with parenthesis. Values are means ± SD. *P < 0.05, **P < 0.01, ***P < 0.001, ****P < 0.0001 and ns, non-significant.

**Fig. S7.**
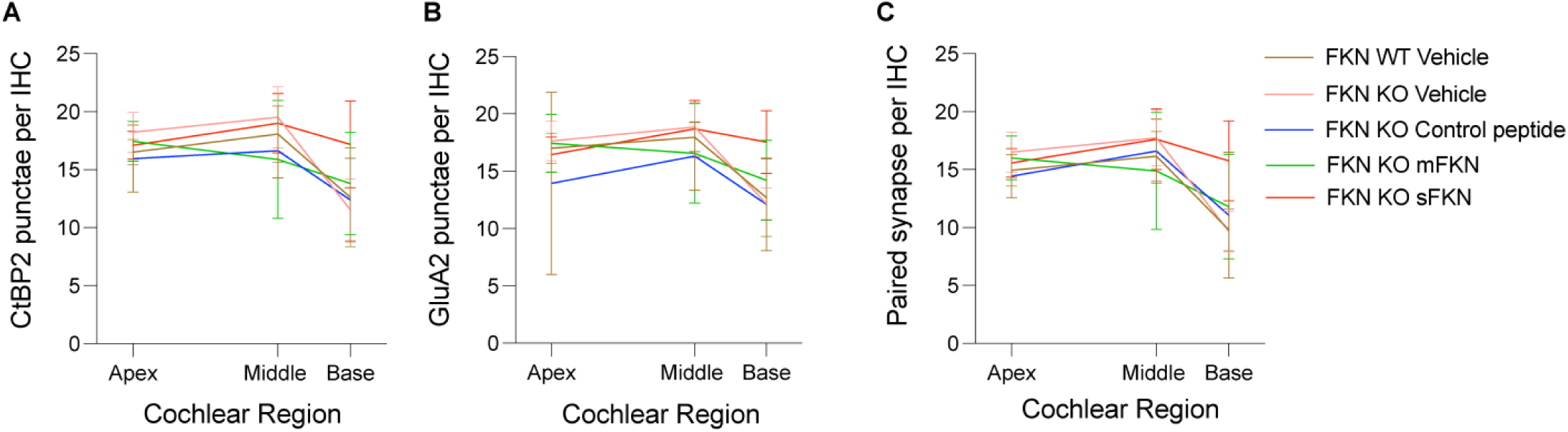
Pre- and postsynaptic punctae and paired synapses per IHC along the cochlear length at 15 DPNE. (A) Absolute CtBP2 punctae per IHC, (B) Absolute GluA2 punctae per IHC and (C) Paired synapse density per IHC at apex, middle and base cochlear regions. Values are mean ± SD. N= 5-9 mice per experimental group.

**Fig. S8.**
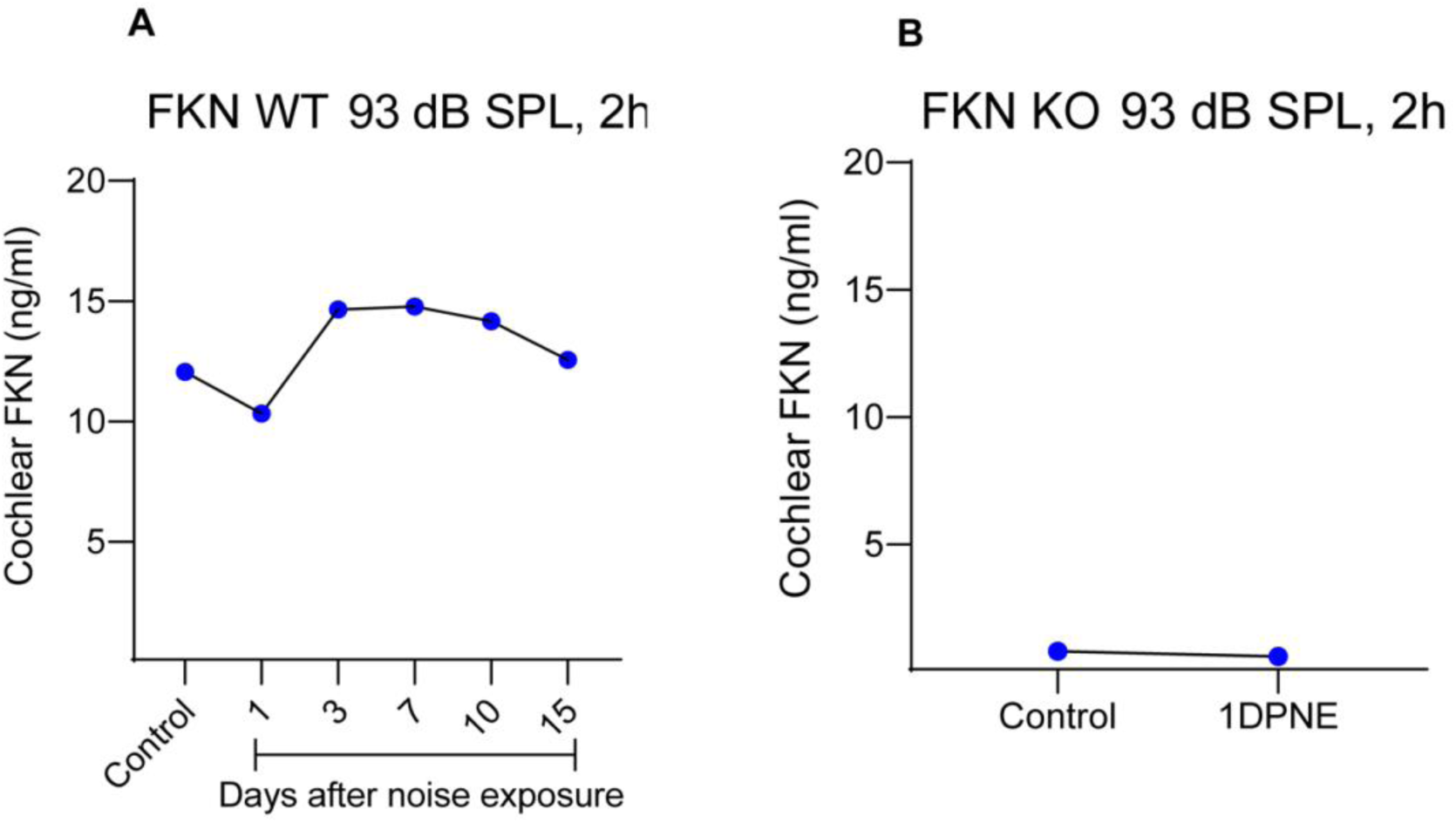
Cochlear endogenous soluble FKN levels after NICS. ELISA based measurement of cochlear endogenous soluble FKN levels in (A) FKN WT mice and (B) FKN KO mice at different time points after NICS. Values are plotted as mean. Experiment was performed once. N=6 cochleae pooled from 3 different mice per time point per genotype to prepare protein lysates and each sample was run in triplicate. DPNE, days post noise exposure.

**Fig. S9.**
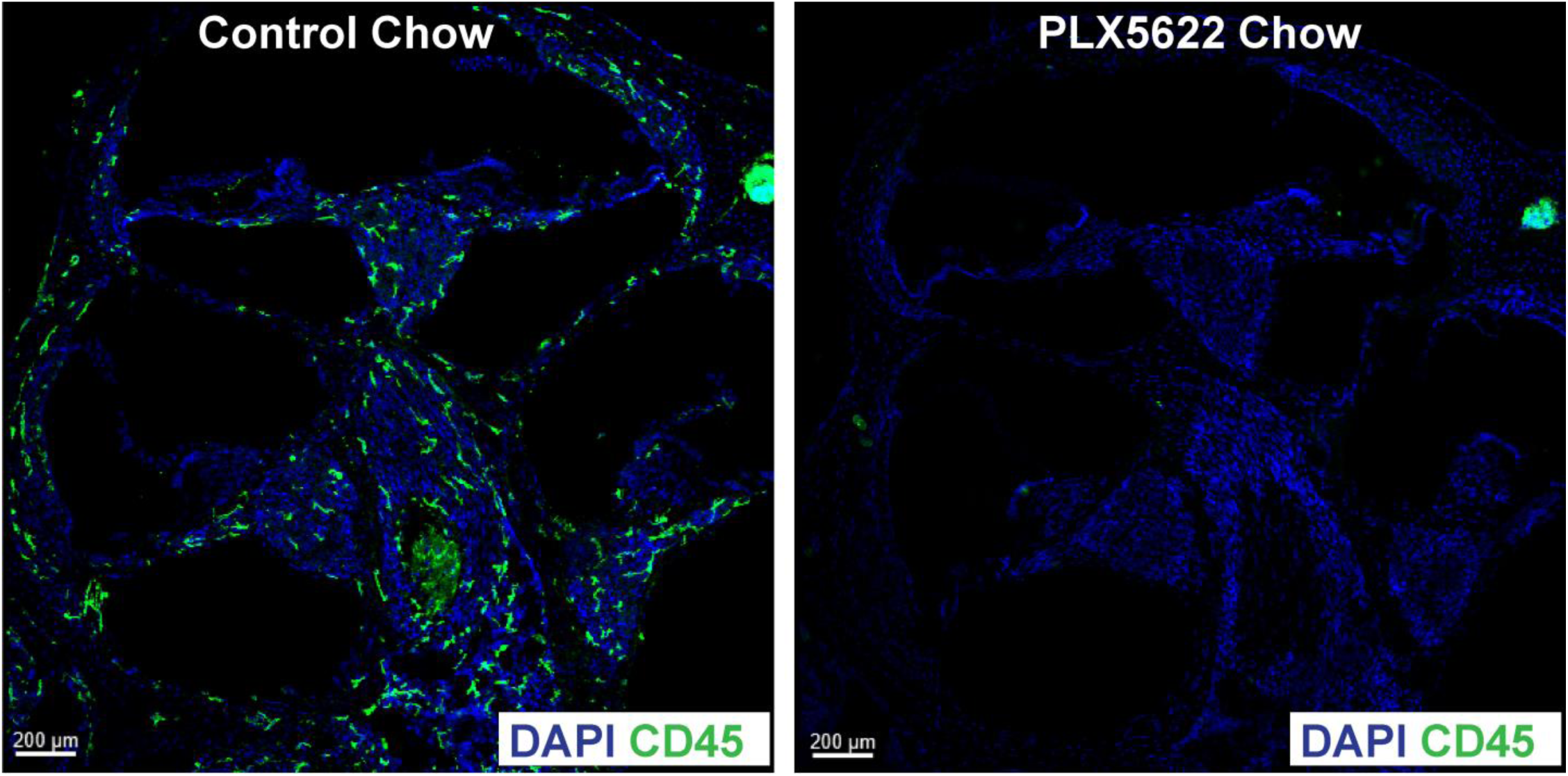
PLX5622-induced depletion of cochlear resident macrophages. Representative images of cochlear mid-modiolar cross-sections from mature FKN WT mice fed on PLX5622 chow for 15 days show a robust depletion of cochlear resident macrophages (green, CD45 leukocytes) when compared to FKN WT mice fed on control chow without the PLX5622 compound for 15 days. Data is from same mice as shown in the main Fig. 5C-F. Also see reference number 13.

**Fig. S10.**
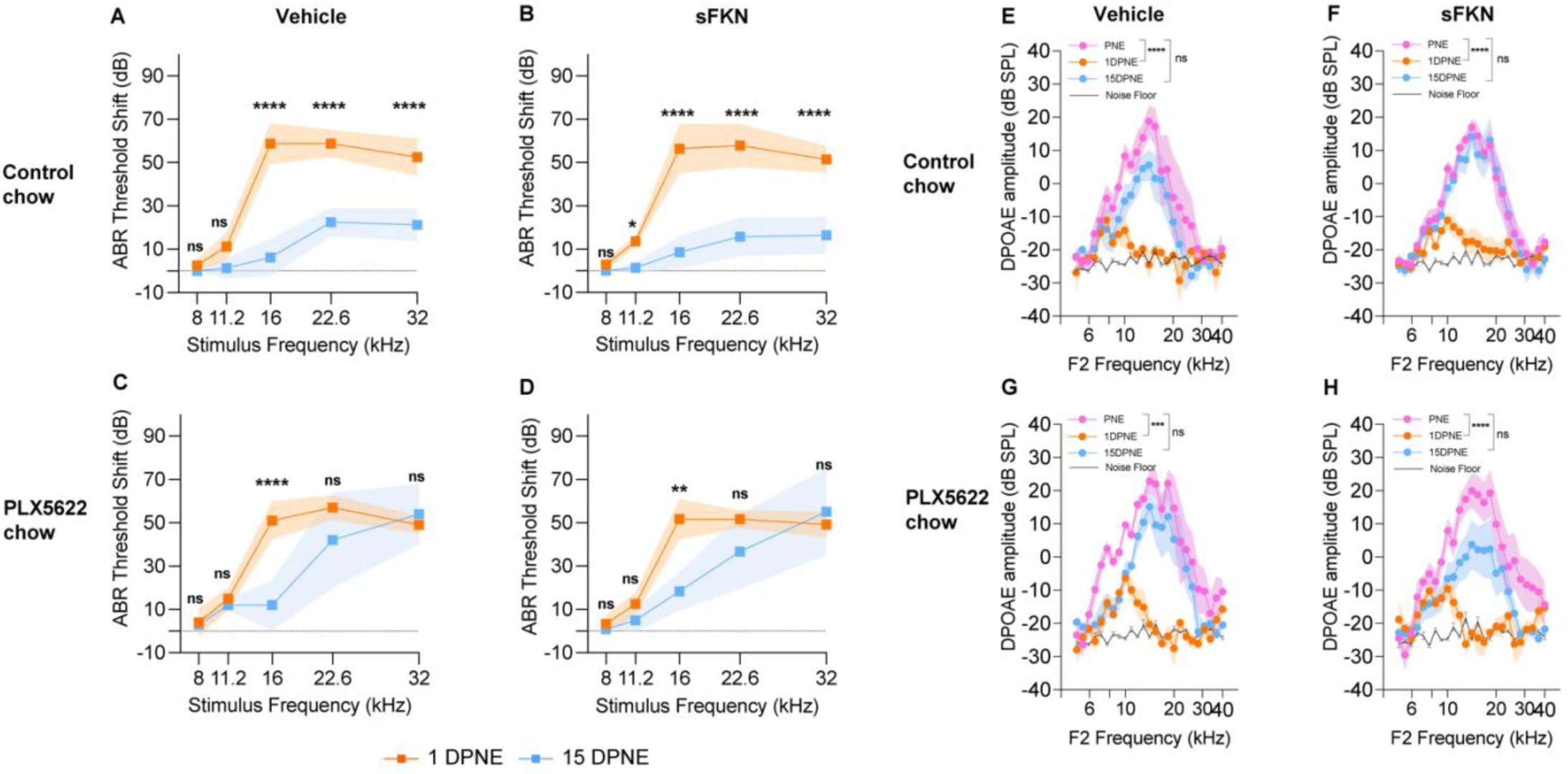
ABR thresholds shifts (A-D) and DPOEA levels (E-H) in vehicle- and sFKN-treated FKN WT mice in the presence (control chow) or absence (PLX5622 chow) of cochlear resident macrophages. N=5-7 mice per experimental group. Values are means ± SD. *Represents comparison between 1DPNE and 15 DPNE (A-D). *Represents comparison between PNE and 1DPNE or 15 DPNE as shown with parenthesis (E-H). 2-way ANOVA, Sidak’s multiple comparisons. *P < 0.05, **P < 0.01, ***P < 0.001, ****P < 0.0001 and ns, non-significant. Dashed line represent threshold shifts prior to noise exposure (baseline, A-D).

**Fig. S11.**
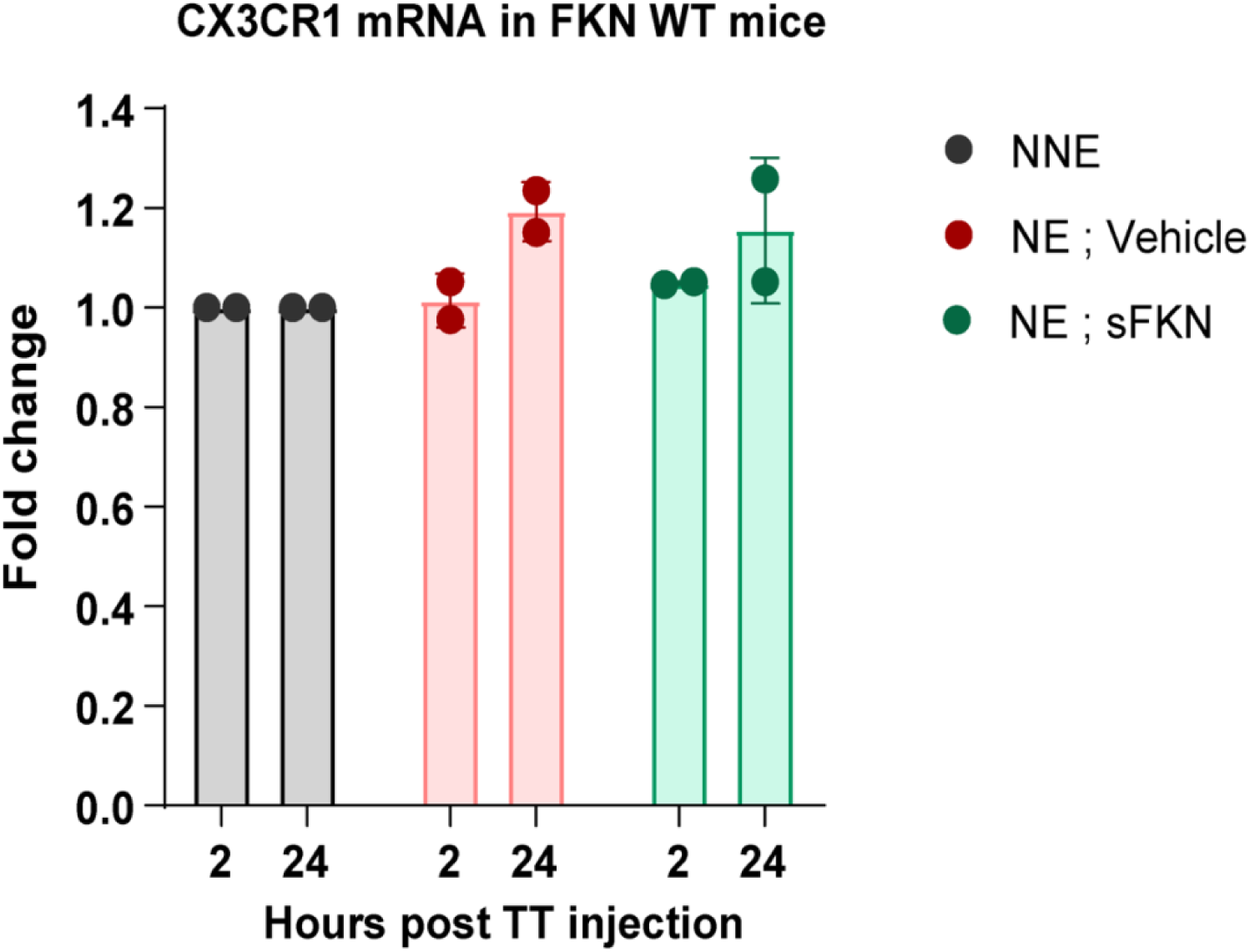
mRNA levels of cochlear CX_3_CR1 receptor. CX_3_CR1 receptor mRNA measured by RT-qPCR in unexposed and noise exposed FKN WT mice after 2 and 24 hours of sFKN peptide TT injection. Values are mean ± SD. N=2 biological replicates per experimental group with 3 cochleae pooled from 3 different mice per group per biological replicate.

**Fig. S12.**
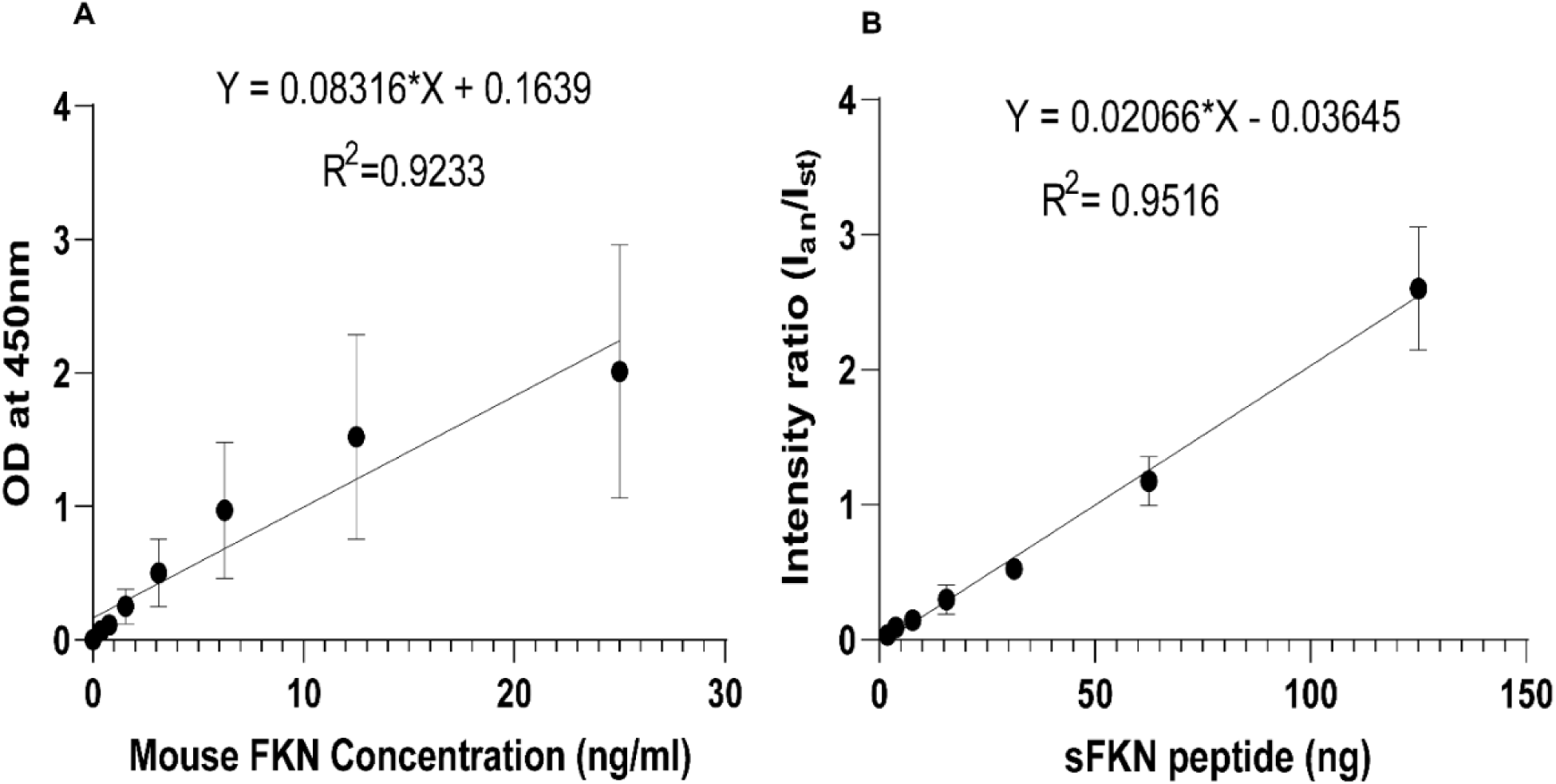
Standard curves for sFKN peptide for (A) ELISA and (B) MALDI-TOF-MS to detect the concentration of transtympanically injected sFKN peptide in the cochlea.

**Fig. S13.**
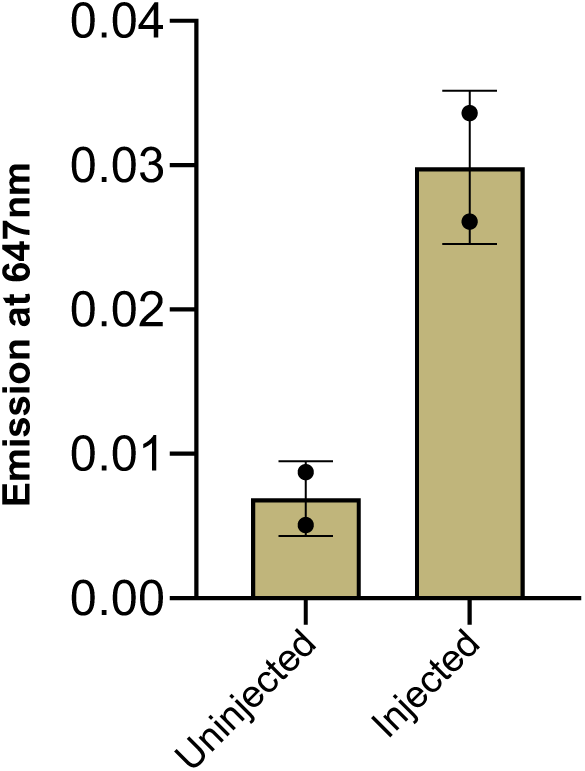
Mean fluorescence intensity of sFKN-Alexa Fluor 647 in cochlear perilymph at 3 hours after TT injection. Values are mean ± SD. N=2 FKN WT mice per experimental group.

## Acknowledgments

We are thankful to Rene Vielman Quevedo for providing training with transtympanic injection in mice, to the translational hearing center auditory and vestibular technology (AVT) core and to the hearing research group at Creighton University for constructive criticism that tremendously improved the manuscript.

## Author contributions

Conceptualization: TK

Methodology: VPM, SPM, DYG, ARS, LB, SVM, TK

Investigation: VPM, SPM, DYG, ARS, LB, SVM, TK Visualization: VPM, SPM, DYG, ARS, LB, SVM, TK

Funding acquisition: TK, VPM

Project administration: TK

Supervision: TK

Writing – original draft: VPM

Writing – review & editing: TK, AC

## Data and materials availability

All data are available in the main text or the supplementary materials.

